# A human-specific regulatory mechanism revealed in a preimplantation model

**DOI:** 10.1101/2025.05.10.653263

**Authors:** Raquel Fueyo, Sicong Wang, Olivia J. Crocker, Tomek Swigut, Hiromitsu Nakauchi, Joanna Wysocka

## Abstract

Stem cell-based human embryo models offer a unique opportunity for functional studies of the human-specific features of development. Here, we genetically and epigenetically manipulate human blastoids, a 3D embryo model of the blastocyst, to investigate the functional impact of HERVK LTR5Hs, a hominoid-specific endogenous retrovirus, on preimplantation development. We uncover a pervasive cis-regulatory contribution of LTR5Hs elements to the hominoid-specific diversification of the blastoids’ epiblast transcriptome. Many of the nearly 700 LTR5Hs genomic insertions in the human genome are unique to our own species. We show that at least one such human-specific LTR5Hs element is essential for the blastoid-forming potential via enhancing expression of the primate-specific *ZNF729* gene, encoding a KRAB zinc finger protein. ZNF729 binds G/C-rich sequences, extremely abundant at gene promoters associated with basic cellular functions, such cell proliferation and metabolism. Surprisingly, despite mediating recruitment of TRIM28, at many of these promoters ZNF729 acts as a transcriptional activator. Together, our results illustrate how recently emerged transposable elements and genes can confer developmentally essential functions in humans.

## Introduction

Nearly half of the human genome is comprised of sequences derived from transposable elements (TEs)^1,2^. While the majority of TEs have lost the ability to transpose, they can significantly impact host gene expression by contributing to various cis-regulatory functions or by giving rise to novel regulatory genes, including transcription factors and non-coding RNAs (reviewed in ^3^). One class of TEs frequently coopted for cis-regulatory functions are endogenous retroviruses (ERVs), also called LTR (Long Terminal Repeat) retrotransposons. ERVs are remnants of ancient retroviral infections of the germline that begun to transmit from parents to offspring following integration into the host genome and, over generations, became fixed in the population^4^. In humans, ERV sequences comprise ∼8.9%^1^ of the genome and originated from retroviruses that invaded the ancestral genome at various points after the divergence of primates from other mammals^5^. Due to their clade- or species-specificity and regulatory potential, LTR retrotransposons have contributed to primate-specific diversification of transcriptomes. Indeed, cis-regulatory elements newly emerged during primate evolution are largely derived from TEs^6,7^.

To propagate and be vertically transmitted, retroviruses had to successfully support their own transcription upon infection of germ cells or pluripotent cells prior to germ cell specification. Thus, the retroviral LTR promoters must have entered the host genome able to engage the pre-existing transcriptional machinery in these embryonic cell types. This feature, along with the reduced DNA methylation during early development^8^, can potentially account for the widespread cooption of ERVs for cis-regulatory functions observed in mammalian preimplantation development, a period of embryogenesis that spans the time from the fertilization of the oocyte to the attachment of the blastocyst to the uterine wall. For example, during mouse preimplantation development, many LTRs function as stage-specific promoters. This has been particularly well-documented for MERVL elements driving the 2-cell stage embryo gene expression program^9–11^ and for the MT2B2 retrotransposon, an element that activates a preimplantation-specific isoform of *Cdk2ap1* essential for embryo development^12^. Stage-specific activation of LTRs has also been documented during human preimplantation^3,13–16^. We previously showed that ERVs of the HERVK (HML-2) family —specifically those carrying LTRs of the LTR5Hs subtype— are transcriptionally activated in human embryos, following embryonic genome activation (EGA) at the 8-cell stage^15^ (Figure 1A). These elements remain active throughout preimplantation development and in the epiblast cells of human blastocysts, where HERVK-encoded proteins are readily detectable^15^. HERVK LTR5Hs is also active in human teratocarcinoma and naive embryonic stem cells, an observation which we and others leveraged to demonstrate that LTR5Hs elements function as long-range enhancers in these cell types, albeit without investigating the functional consequences of such cis-regulatory function^17,18^. HERVK LTR5Hs is the evolutionarily most recent human ERV. It first invaded the genome ∼20 million years ago, after the split of hominoids (apes) from Old World monkeys, and it continued to be active after the split of humans and chimpanzees^19^. As a result of this recent activity, the ∼700 LTR5Hs insertions present throughout the human genome are unique to hominoids, with some being specific to humans^19,20^. The functional impact of HERVK LTR5Hs on preimplantation development, and how this hominoid-specific retrotransposon may have contributed to the transcriptomic and phenotypic divergence of early embryogenesis in humans remains poorly understood.

**Figure 1.**
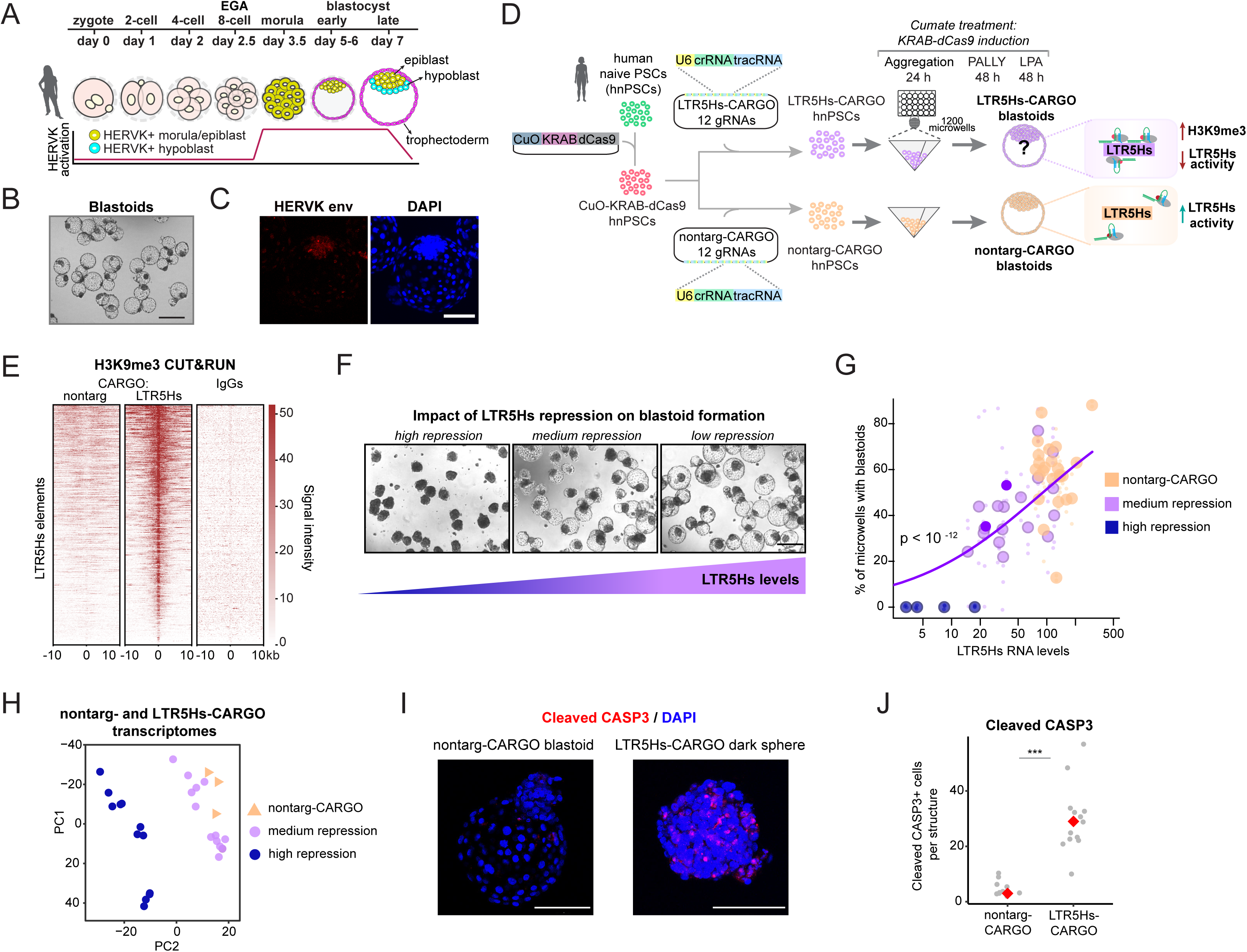
LTR5Hs activity contributes to blastoid formation potential in hnPSC. a. Cartoon depicting stages of human preimplantation development. HERVK expression levels display data in15,32 and are represented in the bottom graph by a red line. HERVK is active in the epiblast (yellow cells) and the hypoblast (cyan cells). EGA: embryonic genome activation. b. Representative bright field image of blastoids generated from wild type hnPSCs, n=3, black bar represents 400 um. c. Representative confocal images of a wild type blastoid immunostained with an antibody against HERVK envelope protein and DAPI for nuclear staining, n=4, white bar represents 100 um. d. Schematics of nontarg-CARGO and LTR5Hs-CARGO hnPSCs and blastoids generation. A cumate-inducible KRAB-dCas9 transgene and a 12-mer CARGO array targeting LTR5Hs elements or a control array (nontarg-CARGO) were integrated into the genome. During the first 24 h, while cells are undergoing aggregation in plates containing 1200 microwells, the cumate treatment is started. Aggregation is followed by 96 h of culture in PALLY and LPA media, blastoid-like structures or aggregates are collected for analysis. Right cartoon summarizes how targeting of LTR5Hs elements by KRAB-dCas9 promotes H3K9me3 deposition and LTR5Hs repression. e. LTR5Hs-CARGO results in localized H3K9me3 deposition across LTR5Hs insertions. Heatmap displaying H3K9me3 CUT&RUN signal over LTR5Hs elements in the human genome in nontarg-, LTR5Hs-CARGO cells, and in the IgGs negative control. f. Bright field images of the structures collected upon LTR5Hs high repression (left, dark spheres), medium repression (middle, dark spheres / blastoid-like structures), and right, no LTR5Hs repression (blastoid-like structures). Black bar represents 400 um. g. Blastoid-forming potential of hnPSC is dependent on LTR5Hs activity (*** p<10-12, beta-regression). Blastoid formation was assessed in 23 LTR5Hs-CARGO (n=3) and 24 nontarg-CARGO (n=2) clonal cell lines. Plotted is proportion (%) of blastoids per well (abscissa) versus LTR5Hs RNA levels (ordinate) as measured by Taqman assay against LTR5Hs elements and normalized to RPL13A RNA levels. Purple line represents LTR5Hs-CARGO regression line. Dark purple circles indicate clones used for scRNA-seq experiments in figures 2 and 3. h. Principal component analysis of the bulk RNA-seq transcriptomes obtained from nontarg-CARGO and LTR5Hs-CARGO clonal cell lines with high or medium expression levels (total 11 clonal cell lines in biological duplicates or triplicates). i. Representative images of blastoid (left, n=11, from three biological replicates) or dark sphere (right, n=13, from three biological replicates) immunostained with the apoptotic marker cleaved-CASP3 (red) and DAPI (blue). White bar represents 100 um. j. Quantification of cleaved-CASP3 immunostainings described in I, unpaired two-tailed test, ***<0.001.

Although the general principles of early development are deeply conserved across mammals^21–23^, many aspects have diverged between species such as humans and mice^22–24^. These observations, along with the fact that ethical and practical limitations largely preclude functional studies in humans^25^, underscore the importance of developing genetically accessible, scalable and ethical models of human preimplantation development. Recent groundbreaking work from many labs established stem cell-based 3D models named blastoids, which recapitulate morphology and formation of the three lineages of the human blastocyst^26–30^. While not without limitations, human blastoids offer unprecedented opportunities to study molecular mechanisms governing species-specific features of human preimplantation development. Here, we epigenetically and genetically perturb HERVK LTR5Hs function in human blastoids. *En masse* manipulation of HERVK LTR5Hs elements reveals their dose-dependent impact on blastoid formation potential and gene regulation. At the single-locus level, we uncover a human-specific LTR5Hs insertion that enhances expression of a gene encoding the zinc finger transcription factor *ZNF729*, promotes proliferation of human naïve pluripotent stem cells (hnPSCs), and is essential for blastoid formation. Although ZNF729 was previously reported as a marker of human naïve and formative pluripotency^31^, its molecular function remained unexplored. We show that ZNF729 binds and regulates G/C-rich promoters of genes involved in fundamental cellular functions. Altogether, our work reveals an evolutionary novel mechanism regulating conserved cellular processes and paves the way for systematic interrogation of TEs or ‘genomic dark matter’ during human embryogenesis.

## Results

### LTR5Hs activity contributes to the blastoid formation potential of hnPSCs

To probe the phenotypic impact of HERVK LTR5Hs activity on human preimplantation development, we turned to a human blastoid model^26^ (Figure 1B). A blastoid is composed of cell types that are analogous to the three cell lineages present in the blastocyst: the epiblast that during natural development gives rise to the embryo proper, the trophectoderm that generates the placenta, and the hypoblast that develops into the yolk sac^21^. Within this framework, we first reanalyzed a comprehensive human embryo single cell (sc)RNA-seq dataset^32^ that became available after we initially reported HERVK expression in the epiblast of human blastocysts^15^. As expected, we observed high HERVK expression in the epiblast lineage and low in the trophectoderm. Interestingly, hypoblast cells also express HERVK (Extended Data Figure 1A and 1C). To verify that blastoids recapitulate the HERVK expression pattern seen in human blastocysts, we reanalyzed published scRNA-seq data from human blastoids^26^. Indeed, this analysis showed that HERVK is expressed in epiblast and hypoblast lineages of human blastoids, as seen in the blastocysts (Extended Data Figure 1B and 1D).

We implemented the blastoid generation protocol^26,33^ using hnPSCs generated from peripheral blood cells by overexpressing NANOG and KLF2^34,35^. The hnPSCs formed blastoids (Figure 1B, Movie 1) with an efficiency of ∼70% (Extended Data Figure 1E), likely an underestimation, as some aggregates/blastoids are accidentally aspirated during the media changes. We benchmarked the blastoid protocol using immunostainings with markers of the three blastocyst lineages (KLF17, NANOG, SUSD2, and IFI16 for epiblast; GATA3 for trophectoderm; SOX17 and GATA4 for hypoblast, Extended Data Figure 1F) and HERVK envelope (Figure 1C), all of which showed staining patterns consistent with those seen in human blastocysts^15,36–38^. Further analysis by scRNA-seq demonstrated that cells dissociated from the blastoids we generated have transcriptome profiles similar to those of the cells originally reported by Kagawa et al.^26^ with analogs of the three blastocyst lineages present (Extended Data Figure 1G).

We previously reported that combining CARGO (Chimeric Array of guide RNA Oligonucleotides)^39^ with CRISPR interference (CRISPRi) allows for efficient and selective *en masse* perturbation of HERVK LTR5Hs function across the genome^17^. Briefly, a multiplexed 12-mer guide RNA (gRNA) array was designed to target the majority of 697 LTR5Hs instances in the genome (hereafter referred to as LTR5Hs-CARGO), along with a control non-targeting array that does not pair anywhere in the human genome (hereafter referred to as nontarg-CARGO). We leveraged this validated system to study the functional impact of HERVK LTR5Hs repression on human blastoid formation. To do this, we generated hnPSCs expressing cumate-inducible catalytically dead version of Cas9 (dCas9) fused to the transcriptional repressor KRAB (dCas9-KRAB) and then introduced LTR5Hs-CARGO or nontarg-CARGO arrays to generate clonal cell lines (Figure 1D). We confirmed that induction of dCas9-KRAB in hnPSCs expressing the LTR5Hs-CARGO but not the nontarg-CARGO resulted in the repression of LTR5Hs-originating transcripts in hnPSCs (Extended Data Figure 2A) and in H3K9me3 deposition across the majority of the LTR5Hs instances in the genome (Figure 1E; see Extended Data Figure 2B for an example of H3K9me3 CUT&RUN genome browser tracks at an individual LTR5Hs instance).

We next asked how LTR5Hs repression affects blastoid-forming potential of hnPSCs. To this end, we induced blastoid generation from 24 distinct nontarg-CARGO and 23 distinct LTR5Hs-CARGO hnPSCs clonal cell lines and concomitantly with cell aggregation, induced dCas9-KRAB expression. We then measured blastoid formation efficiency as a function of LTR5Hs expression levels in these clonal lines (Figure 1D, F and G). We note that some of the LTR5Hs-CARGO lines showed nearly complete LTR5Hs repression (hereafter called ‘high repression’ clones), whereas others showed medium or low repression levels. We observed a correlation between LTR5Hs expression and blastoid-forming potential, with near-complete repression of LTR5Hs activity being incompatible with blastoid formation and instead resulting in structures resembling dark spheres (beta regression, p-value <10^-12^). At the intermediate level of repression, blastoid-like structures still formed, albeit at reduced efficiencies, whereas hnPSCs lines with poor LTR5Hs repression formed blastoids at efficiencies comparable to the nontarg-CARGO control cell lines (with some variability, likely attributable to clonal effects). To confirm that defects in blastoid formation are not due to LTR5Hs-CARGO off-target effects, we cloned a new array of guide RNAs, fully orthogonal to our original LTR5Hs-CARGO array (named LTR5Hs-Ortho-CARGO). We integrated this array into the hnPSCs genome together with a cumate inducible dCas9-KRAB and confirmed that it drove efficient repression of LTR5Hs and its selected gene targets (Extended Data Figure 2C). Importantly, in agreement with our LTR5Hs-CARGO result, LTR5Hs-Ortho-CARGO hnPSCs also failed to form blastoids (Extended Data Figure 2D).

The retrotransposon HERVK retains proviral copies with coding capacity for viral proteins and these can be detected in human embryos^15^. To explore the idea of the viral proteins being responsible for the failed blastoid formation phenotype, we performed rescue experiments by genomic integration of multiple copies of a constitutively active transgene encoding the HERVK viral proteins gag, pro, and pol^40^. This transgene failed to restore the blastoid formation capacity of the LTR5Hs-CARGO high repression hnPSCs, suggesting that HERVK viral proteins alone are not responsible for the dark spheres phenotype (Extended Data Figure 2E). Together, these results indicate that LTR5Hs activity affects the blastoid-forming potential and suggest a non-neutral contribution of this hominoid-specific transposon to human preimplantation development.

### Dose-dependent gene expression changes upon LTR5Hs-repression in hnPSCs

The different capacity of LTR5Hs-CARGO clones to generate blastoids prompted us to analyze gene expression changes in high and medium repression hnPSCs following dCas9-KRAB induction for 96 h (we note that hnPSCs cannot be maintained long-term after LTR5Hs repression). We performed bulk RNA-seq in LTR5Hs-repressed and nontargeting hnPSCs (a total of 4 high repression and 5 medium repression LTR5Hs-CARGO clonal cell lines and two nontarg-CARGO clones in biological duplicates or triplicates). Principal component analysis (PCA) revealed that while the LTR5Hs-CARGO clones separated from the control nontarg-CARGO clones, those with medium repression levels were clustering much closer to the controls, while those with the high repression were more distant (Figure 1H). In agreement, differential gene expression analysis confirmed stronger misregulation of gene expression in high vs medium repression clones relative to the nontarg-CARGO controls, in both number of affected genes and magnitude of the observed effects (Extended Data Figure 2F and G). Of note, gene ontology analysis of transcripts dysregulated in the high repression clones but not in the medium repression clones revealed categories related to embryo morphogenesis and immune response among others, altogether suggesting that the high repressing clones undergo an additional level of gene dysregulation (Extended Data Table 1).

Next, we sought out to investigate the dark spheres phenotype obtained upon near-full LTR5Hs-repression. These aggregates of cells do not show signs of cavitation and look homogeneous under bright field (Figure 1F, left panel). To analyze the nature of these structures, we performed bulk RNA-seq and compared it to bulk RNA-seq of control blastoids. We identified differentially expressed genes and detected a clear separation of these transcriptomes in the PCA space (Extended Data Figure 2H, and I). Gene ontology analysis of the differentially expressed genes revealed categories related to morphogenesis, migration, cell proliferation, Wnt pathway, immune response and others (Extended Data Table 1). Among the upregulated genes, we also detected genes pointing to apoptosis (e.g. *CASP7*) and this prompted us to systematically investigate if apoptotic genes are differentially regulated. Indeed, comparison of the differentially regulated genes with a set of curated apoptosis genes from GSEA^41^ identified several transcripts upregulated in the dark spheres which are typical of apoptotic cells (Extended Data Figure 2J). To confirm this result, we stained blastoids and dark spheres with the apoptotic marker Cleaved-CASP3. While blastoids displayed a median of three Cleaved-CASP3+ cells, for dark spheres the median was 29, almost 10 times higher. This result confirms that under high LTR5Hs repression conditions, the hnPSC undergo widespread gene expression changes incompatible with blastoid formation and consistent with an apoptotic phenotype.

### Loss of LTR5Hs activity impacts lineage identity in human blastoids

Finding that hnPSCs lines with medium level of LTR5Hs repression still retain ability to form blastoid-like structures, albeit with lower efficiency, offered us an opportunity to address whether these hypomorphic blastoids show lineage defects. To this end, we induced these medium repression LTR5Hs-CARGO clonal cell lines to form blastoids in parallel with nontarg-CARGO hnPSC lines and performed immunostainings using known markers of the preimplantation epiblast (KLF17, SUSD2^32,36,42^), the hypoblast (GATA4^32,38^), and the trophectoderm (GATA3^32,37^). We observed that LTR5Hs-CARGO blastoids had a diminished number of cells marked by KLF17 and GATA4, and an overall decreased signal of SUSD2, suggesting defects in the epiblast and hypoblast lineages (Figure 2A, 2B, and Extended Data Figure 3A). Of note, this decrease in the number of KLF17+ or SUSD2+ cells was not due to a loss of blastoids’ epiblast cells, as the general epiblast marker NANOG is expressed both in nontarg-CARGO and LTR5Hs-CARGO blastoids (Extended Data Figure 3B). In contrast, we detected an increased number of GATA3-positive cells, and this was accompanied by smaller inner cell mass (ICM) to trophectoderm ratios, consistent with an expansion of the GATA3-positive trophectoderm compartment (Figure 2A, 2B, and Extended Data Figure 3C).

**Figure 2.**
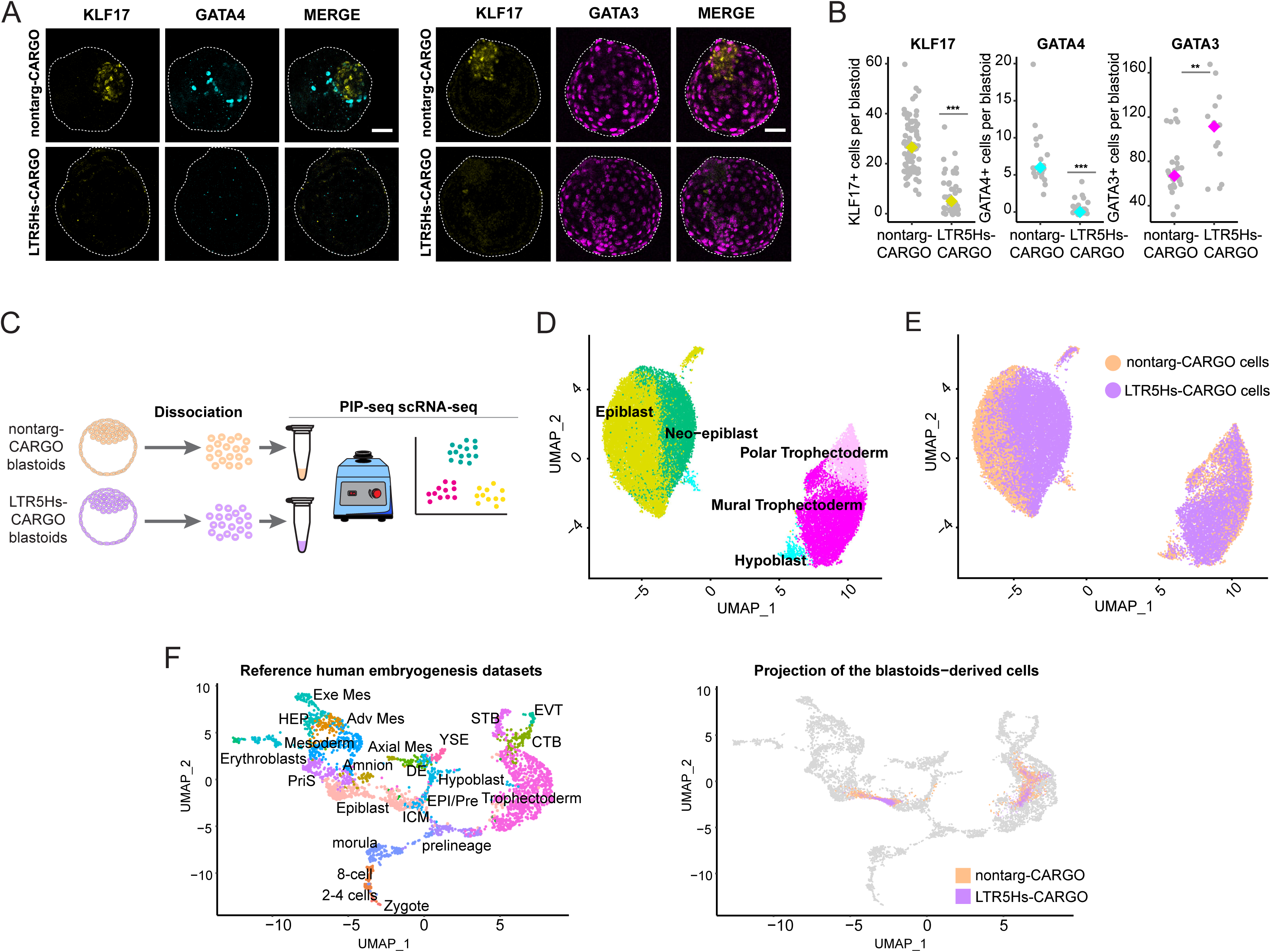
LTR5Hs activity is required for proper lineage acquisition in blastoids. a. Representative confocal images of nontarg-CARGO and LTR5Hs-CARGO blastoids stained with lineage-specific antibodies (yellow: KLF17, epiblast marker, nontarg-CARGO n=78, LTR5Hs-CARGO n=41; cyan: GATA4, hypoblast marker, nontarg-CARGO n=22, LTR5Hs-CARGO n=24; magenta: GATA3 trophectoderm marker, nontarg-CARGO n=27, LTR5Hs-CARGO n=14). Stained blastoids represent at least four biological replicates. White bar represents 50 um. b. Counting of cells showing positive staining for the indicated markers in the nontarg-CARGO and LTR5Hs-CARGO blastoids detailed in a. Grey dots represent the number of cells in individual blastoids. Yellow, cyan, or magenta dots repre-sent median. Unpaired two-tailed test, ***<0.001, **<0.01. c. Schematics of Particle-templated instant partition sequencing (PIP-seq) of single cells obtained from nontarg- or LTR5Hs-CARGO blastoid dissociation. d. and e. Uniform manifold and approximation projection (UMAP) of transcriptomes from single cells dissociated from nontarg-CARGO blastoids and LTR5Hs-CARGO blastoids. d. Colors indicate cells belonging to the same lineage-specific clusters. e. Colors indicate the genotype of origin (orange: nontarg-CARGO blastoids and purple: LTR5Hs-CARGO blastoids). f. UMAP of a reference collection of human embryo single cell RNA-seq transcriptomes29,32,45–49 (left) and projection of nontarg-CARGO (orange) or LTR5Hs-CARGO (purple) PIP-seq results in that UMAP (right).

To systematically interrogate changes in gene expression and lineage allocation associated with LTR5Hs repression, we profiled transcriptomes of single cells by PIP-seq (Particle-templated instant partition sequencing)^43^. For this purpose, we selected two medium repression hnPSC lines presenting blastoid formation efficiencies of 44% on average (highlighted as dark purple circles, Figure 1G) and two control nontarg-CARGO lines (Figure 2C). We generated a global embedding of all samples (see Methods) amounting to a total of 31028 cells and 37468 genes and performed cell cluster annotation based on well-established markers of the blastocyst lineages^32^ (Figure 2D, Extended Data Figure 3D). In parallel, we colored cells based on their genotype of origin (i.e. nontarg- or LTR5Hs-CARGO) (Figure 2E). Cells expressing markers of all three blastocyst lineages were recovered in our analysis (Figure 2D, Extended Data Figure 3D). Within the trophectoderm lineage, we could further distinguish two clusters which we assigned as mural and polar trophectoderm based on expression of *NR2F2*, *PGF*, and *CYP19A1* in the latter^32,44,45^ (Extended Data Figure 3D). Projection of the scRNA-seq transcriptomes into a collection of human embryo datasets^29,32,45–48^ using a prediction tool^49^ confirmed that the nontarg-CARGO blastoids dissociated cells matched transcriptomes of the preimplantation embryo, with cells projected into the epiblast, the hypoblast and the trophectoderm (Figure 2F, colored in orange). Additionally, increased resolution (from 0.2 to 1) of clustering failed to call amnion or mesoderm clusters as was the case in Kagawa et al.^26^. In agreement, amnion markers (*GABRP* or *ISL1*) and mesoderm markers (*APLNR* and *CRABP2*) were either not expressed or did not overlap a specific cluster (Extended Figure 3E).

**Figure 3.**
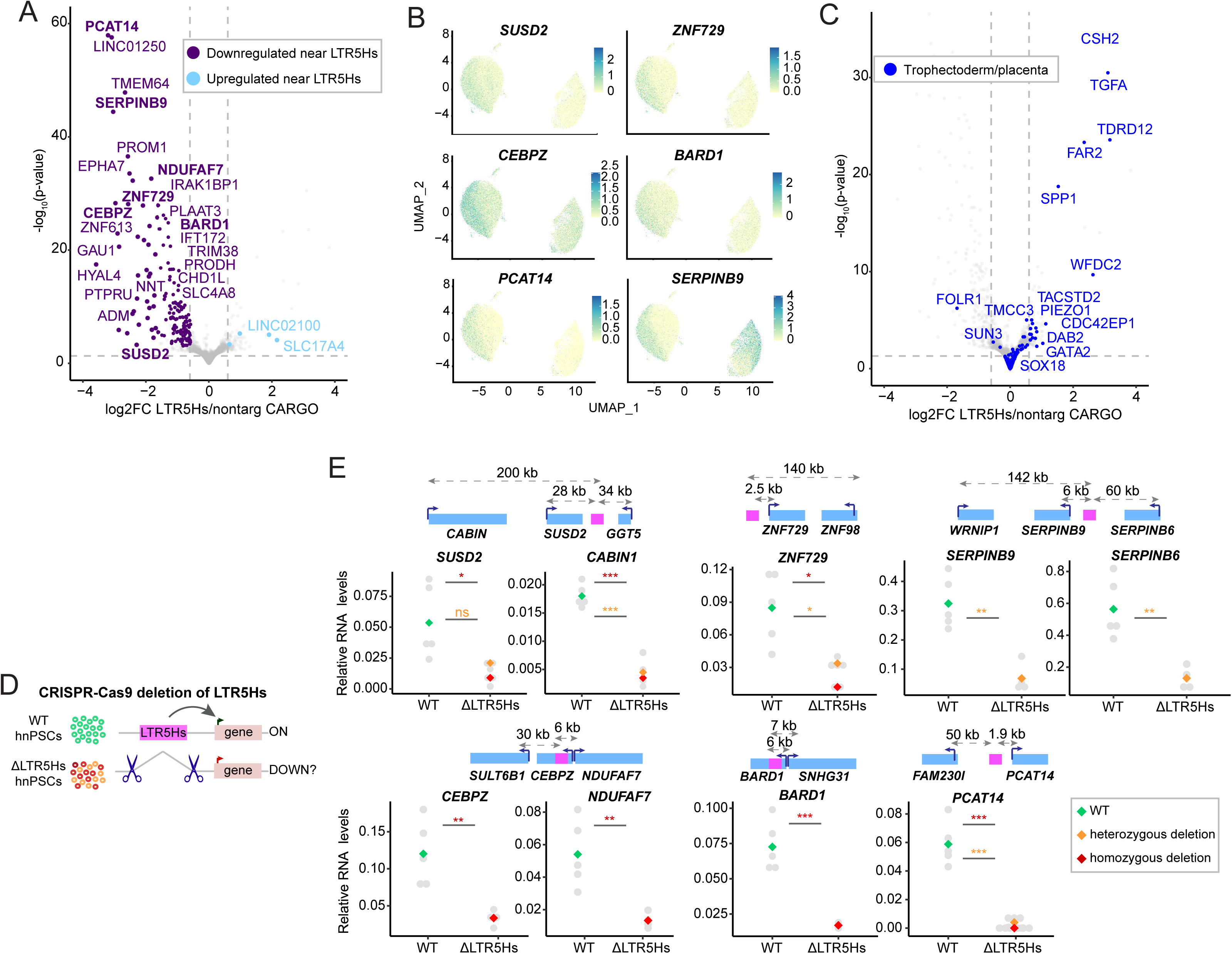
Cis-regulatory activity of LTR5Hs regulates the blastoids’ epiblast transcriptome. a. Volcano plot representing gene expression changes in LTR5Hs-versus nontarg-CARGO blastoids’ epiblast cells (using combined cells from epiblast and neo-epiblast clusters). In a. Purple and light blue represent genes within 250 kb of an LTR5Hs, grey dots represent any other gene. Bold indicates genes further explored in e. Vertical dashed lines indicate fold change < −1.5 or > 1.5 and horizontal dashed line indicates p-value 0.05. b. UMAP of the selected downregulated genes showing higher expression in the blastoids’ epiblast (left cluster) compared to the neo-epiblast cluster (right). c. Volcano plot representing gene expression changes in in LTR5Hs-versus nontarg-CARGO blastoids’ epiblast cells (using combined cells from epiblast and neo-epiblast clusters). Only genes from the trophectoderm or placenta lineages obtained from literature searches and 32,45 are colored in dark blue, the rest of genes are grey color. Vertical dashed lines indicate fold change < −1.5 or > 1.5 and horizontal dashed line indicates p-value 0.05. d. Strategy for testing LTR5Hs-sequence dependency of the observed gene expression changes. Top cartoon shows wild type (WT) hnPSCs containing an LTR5Hs element in proximity of an active gene putatively regulated by such LTR5Hs. Bottom cartoon shows cells displaying homozygous (red) or heterozygous (orange) deletions of the LTR5Hs following CRISPR-Cas9 genome editing. The impact of deletion on the expression of the candidate target gene is tested by RT-qPCR. e. RT-qPCR results of the expression of the indicated genes in wild type or ΔLTR5Hs hnPSCs. RNA values are normalized relative to RPL13A. Above each plot, schematic depiction of the locus is shown. Blue rectangles indicate genes, pink indicates the closest LTR5Hs element, the number on top of the dashed arrows displays distance from the promoter to the LTR5Hs. Grey dots represent expression values obtained in each clone, green, orange, and red dots represent median values. Unpaired two-tailed test, ***<0.001, **<0.01, *<0.05, asterisk color indicates p-value calculated for the homozygous (red) or heterozygous (yellow) clones. ns: not significant.

We observed the emergence of a new epiblast-adjacent cluster, populated mostly by cells from the LTR5Hs-CARGO blastoids (dark green in Figure 2D, purple color in Figures 2E and Extended Data Figure 3F); we termed this cluster ‘neo-epiblast’ by virtue of its transcriptional proximity to the epiblast. While the neo-epiblast cells originated predominantly from the LTR5Hs-CARGO blastoids, the epiblast cluster was populated mostly by cells from the nontarg-CARGO blastoids, suggesting that the new epiblast cluster is a result of gene expression changes caused by the LTR5Hs repression. To examine if this expression shift is associated with change in cell fate, we projected the LTR5Hs-CARGO dissociated cells into the human embryo reference datasets. Interestingly, the LTR5Hs-CARGO epiblast cells clustered with a less mature preimplantation epiblast compared to the nontarg-CARGO epiblast-like cells (Figure 2F, Extended Data Figure 3G), suggesting that LTR5Hs repression affects epiblast maturation in blastoids. Nonetheless, this result should not be understood as HERVK LTR5Hs repression promoting a more pluripotent state, because as revealed by a few diagnostic markers of naïve, formative, and primed pluripotency in the epiblast, the only core naïve pluripotency transcription factors that change are *DPPA5* which displays higher expression and *KLF17* (lower expression). We also observe a minor decrease in the primed pluripotency markers, consistent with the neo-epiblast cells aligning more with a more immature epiblast (Extended Data Figure 4A, 4B, and 4C).

In addition to changes in the epiblast, we observed a reduced allocation of the LTR5Hs-CARGO cells to the hypoblast cluster, marked by *PDGFRA*, *GATA4*, *FOXA2* and other canonical hypoblast genes (Figure 2D, 2E, 2F, and Extended Data Figure 3D). Trophectoderm was the least affected by LTR5Hs repression overall, in agreement with the lowest expression of HERVK in the trophectoderm lineage and our GATA3 immunostaining results. However, in contrast to the immunostainings, we did not detect an increased proportion of the trophectoderm cells in the scRNA-seq analysis of the LTR5Hs-CARGO blastoids. Nonetheless, we do note that scRNA-seq may not be accurate for trophectoderm cell counting, as we have observed an accelerated lysis of trophectoderm cells upon blastoid dissociation prior to cell capture. In agreement with such possibility, the overall proportion of trophectoderm cells was systematically lower in our scRNA-seq analyses compared to immunostainings, irrespective of the LTR5Hs activity status. Beyond this caveat, we noticed diminished contribution of the LTR5Hs-CARGO cells to the polar trophectoderm cluster and a decreased expression of IL6, a highly expressed interleukin in the polar trophectoderm that signals to the epiblast (Figure 2D-E and Extended Data Figure 4D). Interestingly, maturation of the polar trophectoderm is dependent on the signals from the epiblast^26^, raising a possibility that this effect may be an indirect consequence of the defective epiblast.

To test if these lineage defects were recapitulated in 2D hnPSCs cultures, we analyzed the impact of LTR5Hs repression on direct trophectoderm and hypoblast differentiations, utilizing well established protocols^50,51^ and performing quantifications with flow cytometry. LTR5Hs repression did not change the potential of hnPSCs to differentiate into trophectoderm cells (TROP2+ cells) (Extended Data Figure 4E, quantified in 4F) or hypoblast (ANPEP+ cells), if anything the hypoblast differentiation efficiency was slightly elevated in the highly repressed clones (Extended Data Figure 4G, quantified in 4H). We postulate that this seemingly disparate result stems from differences between the 2D hypoblast differentiation and blastoid formation protocols, whereby the former includes 7 cytokines to promote hypoblast differentiation, and the latter relies uniquely on signals from the epiblast. Thus, the hypoblast defect we observe in blastoids may be an indirect consequence of perturbed epiblast function or maturation. Overall, our results show that LTR5Hs repression disrupts lineage determination and allocation within human blastoids and results in a major change in the blastoids’ epiblast transcriptome.

### LTR5Hs-mediated regulation of gene expression in the blastoids’ epiblast

To systematically investigate transcriptome changes occurring in the blastoids’ epiblast-like cells upon LTR5Hs repression, we performed differential gene expression analysis of our scRNA-seq data, combining cells from the epiblast and neo-epiblast clusters (epiblast-agnostic pseudobulk analysis). We called transcripts differentially expressed in the nontarg-vs the LTR5Hs-CARGO epiblast cells using DESeq2^52^ and identified 255 and 87 transcripts that were downregulated and upregulated, respectively (FDR 5%, fold-change < −1.5 or > 1.5) (Extended Data Figure 5A, Extended Data Table 2). Of note, only 122 and 24 of these genes (35% and 7%) were also significantly changed in the high and medium repression hnPSCs clones respectively, highlighting distinct gene regulatory outputs of LTR5Hs in 2D hnPSCs culture versus the 3D human blastoid model. From the transcripts differentially expressed in the epiblast, we first focused on the downregulated set. Given that in the teratocarcinoma cells LTR5Hs elements function as transcriptional enhancers with long-range effects on gene expression over distances of up to 250 kb from their target gene promoters^17^, we examined which of the downregulated epiblast gene promoters reside within 250 kb of the LTR5Hs insertion. We found that 84% of the protein-coding genes in the downregulated group lie within 250 kb of the LTR5Hs, compared to the 1.62 Mb of median distance to LTR5Hs of all human protein coding genes. This observation suggests direct regulation by LTR5Hs in *cis* (Figure 3A, highlighted in purple). As expected, given that neo-epiblast cluster is mostly populated by LTR5Hs-repressed cells, downregulated genes also display a clear decrease in expression when comparing neo-epiblast and epiblast clusters in the UMAP plots (Figure 3B).

We next looked at the upregulated genes. In contrast to the downregulated genes, they were typically not located near LTR5Hs (only 5% of the upregulated TSS were within 250 kb from the LTR5Hs) (Figure 3A), suggesting indirect effects. Interestingly, however, the upregulated genes showed a clear signature, with the majority of them being associated with trophectoderm and/or placenta development according to previously established markers and literature searches (Figure 3C). As examples, the gene *CSH2*, encodes for the placental hormone named lactogen^53^; GATA2 is a key transcription factor in the trophectoderm lineage^32^; CDC42EP1 participates in trophectoderm sorting and migration^54^; TACSTD2, also known as TROP2, and DAB2 are common trophoblast markers^32,50^; and notably, the transforming growth factor alpha, TGFA, has been described to increase blastocoel size in mouse blastocysts^55^, suggesting it could be contributing to the increase in the number of GATA3+ cells in the LTR5Hs-CARGO blastoids (Figure 2A and 2B and Extended Data Figure 3C). Altogether, these observations are consistent with the LTR5Hs elements possibly regulating blastoids’ epiblast gene expression in *cis*. Perturbation of the epiblast regulatory program is in turn associated with an indirect upregulation of genes involved in the trophectoderm specification and maturation, as the inductive signals for this fate are present in the developing blastoid. Nonetheless, it is important to note that despite this upregulation, these cells remain closer to the naïve pluripotent state and retain expression of key naïve pluripotency genes (Figure 2F, Extended Data Figure 4B).

### LTR5Hs elements are enhancers for human epiblast genes

Enhancers are defined as genetic elements and as such, we sought to address whether the observed effects on the epiblast gene expression are dependent on the LTR5Hs DNA sequence. H3K9me3 deposition upon dCas9-KRAB recruitment is localized and does not spread beyond a few kilobases from the targeted element (see also Extended Data Figure 2B), whereas the transcriptional effects we observe upon LTR5Hs repression occur at much longer ranges. Nonetheless, several relevant downregulated epiblast genes do have LTR5Hs close to the promoter (i.e. < 5kb). Thus, both to establish sequence-dependency and to ensure that our results are not an artifact of ectopic silencing, we selected six different LTR5Hs elements in the vicinity of the downregulated genes whose products have known or potential functions in the epiblast and that show clear enrichment in the blastoids’ epiblast when compared to the neo-epiblast (Figure 3B). For example, SUSD2 is an established marker of human naive pluripotency^31,42,56^, ZNF729 a marker of naïve and formative pluripotency^31^, CEBPZ is a transcription regulator highly expressed in the epiblast, PCAT14 is a long-noncoding RNA implicated in proliferation^57^, BARD1 is a partner of BRCA1 and is essential for embryonic development in mice^58^, whereas SERPINB9 belongs to the serine proteinase inhibitor superfamily and, among other functions, has been related to cancer stem cell self-renewal^59^.

We used CRISPR-Cas9 genome editing to engineer a series of hnPSCs clonal cell lines in which each of the aforementioned six LTR5Hs elements has been deleted, one at a time, either in a homozygous or heterozygous setting, with multiple clonal lines for each element (Figure 3D). We then used reverse transcription coupled to quantitative PCR (RT-qPCR) to measure effects of each deletion on gene expression in comparison to wild type cells that have undergone clonal selection in parallel. Given that the selected epiblast genes are also expressed in naive pluripotency, we conducted our measurements in hnPSCs. For each of the analyzed cell lines, we observed that LTR5Hs deletion was associated with a downregulation of the candidate epiblast target gene (Figure 3E). In the case of *SERPINB9*, homozygous deletions could not be recovered, suggesting that both the LTR5Hs and the target gene itself may be essential for hnPSCs survival. We also noticed a slower growth of the clones in cells with LTR5Hs deletion at the *ZNF729* locus, suggesting a potential role in hnPSCs proliferation. Apart from this, the derived clonal cell lines were morphologically undistinguishable from wild type hnPSCs (Extended Data Figure 5C).

To ensure that these genetic deletions were not giving rise to DNA copy number variations and to identify other potential targets of the deleted LTR5Hs copies, we scanned genomic loci surrounding the six LTR5Hs elements selected for deletion. We observed that these LTR5Hs elements contain other genes in the vicinity, beyond those already examined (Figure 3E and Extended Data Figure 5B, see diagrams on top of each plot). We therefore analyzed expression of these other potential gene targets at the six loci. Interestingly, in some cases LTR5Hs deletion affected expression of more than one gene at the locus (e.g. *SUSD2* and *CABIN1*; *SERPINB9* and *SERPINB6*; *CEBPZ* and *NDUFAF7*) (Figure 3E), whereas in other cases, there was no effect on the other neighboring gene (e.g. *GGT5, ZNF98*, *FAM230I*, *SNHG31, WRNIP1*) (Extended Data Figure 5B), suggesting a degree of selectivity in promoter responsiveness to the LTR5Hs enhancers. For each deletion, we observed at least one nearby gene whose expression remained unaffected, confirming specific deletions that are not broadly altering the locus. Overall, promoters of genes downregulated upon LTR5Hs deletions were located at distances ranging from 1.9 – 200 kb from the LTR5Hs and represented different arrangements in relation to orientation between the LTR5Hs element and the promoter. Considering that dependence on DNA sequence, ability to activate distally to the promoter and independence of orientation are all hallmarks of enhancers^60^, we conclude that LTR5Hs elements likely function as hominoid-specific enhancers for many human epiblast genes.

### LTR5Hs contributed to hominoid-specific diversification of the epiblast transcriptome

Given the prominent enhancer function of LTR5Hs in the blastoids’ epiblast, we hypothesized that this retrotransposon may have played an important role in the hominoid-specific diversification of the epiblast transcriptome. To explore this, we have investigated the levels of expression and conservation of LTR5Hs-regulated genes between humans, marmoset (a primate that lacks HERVK LTR5Hs), and the most well studied mammalian embryo model, the mouse. For this purpose, we first identified candidate direct target genes of the LTR5Hs as those downregulated upon LTR5Hs repression in the epiblast cells of blastoids and located within 250 kb of the LTR5Hs insertion (hereafter referred to as ‘LTR5Hs target genes’). Then, we looked at the conservation of the LTR5Hs target genes and at their expression in the epiblast by drawing upon a previous study comparing the transcriptomes of staged-match human, marmoset, and mouse preimplantation embryos^61^. Out of the 144 LTR5Hs target genes present in the datasets, 37 did not have an ortholog in the mouse, and thus are not expressed in the mouse epiblast (Figure 4A). Using the Gentree database^62^, we assigned these genes to evolutionary branches noticing that at least four genes (specifically, *ZNF729*, *CR1L*, *ZNF676*, and *NBPF12*) are unique to primates, while the rest are evolutionarily older (Extended Data Table 3).

**Figure 4.**
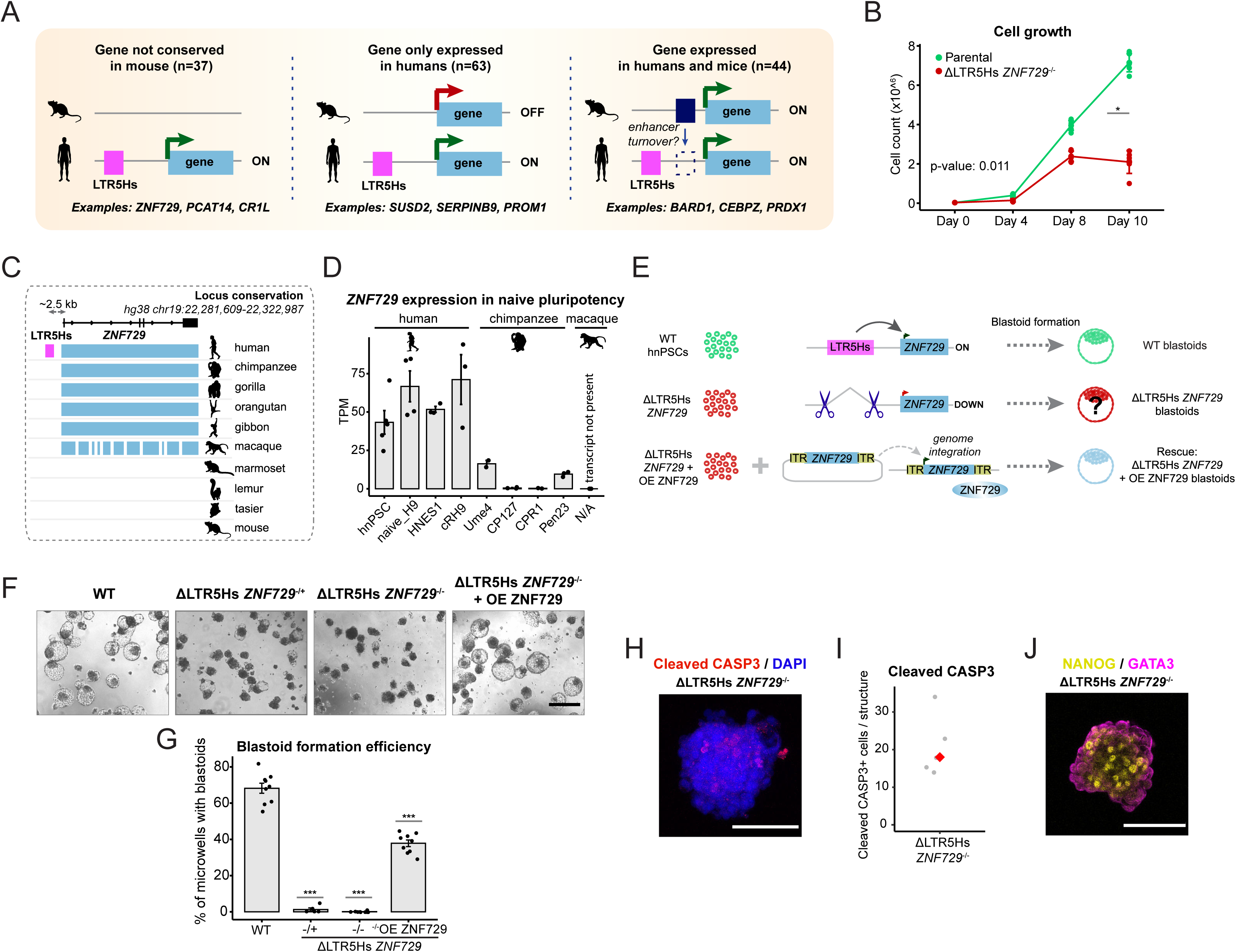
Human-specific LTR5Hs regulation of a primate-specific gene is essential for blastoid formation. a. Expression and conservation of the LTR5Hs-regulated genes in mice. Human epiblast genes regulated by LTR5Hs in cis were divided into three groups, based on the status of expression in the mouse epiblast and conservation of the gene itself using data published in 61. The number of genes in each of the three groups is indicated on top, examples of genes within each group are shown at the bottom. b. Cell proliferation curves in parental (green) and ΔLTR5Hs ZNF729-/-hnPSCs (red), n=3. One-way ANOVA, *<0.05. c. Cartoon displaying the conservation of the primate-specific gene ZNF729 and its regulatory landscape in humans, chimpanzees, gorillas, orangutans, gibbons, rhesus monkeys, marmosets, lemurs, tarsiers, and mice. LTR5Hs element is only present in humans. Data adapted from Cactus UCSC tracks and Gentree databases. d. Expression of ZNF729 in a collection of human50,51,130 and chimp naïve pluripotent stem cells68, rhesus monkeys lack the transcript annotation (Mmul_10, RheMac10). Each dot represents a bulk-RNA-seq replicate. e. Schematic representation of the experimental approach to address the essentiality of the human-specific LTR5Hs element located near the ZNF729 gene in blastoid formation. Top section represents wild type hnPSCs (green), with an intact LTR5Hs element functioning as an enhancer of the ZNF729 gene and which form blastoids. Middle section shows ΔLTR5Hs ZNF729 hnPSCs (red, i.e. cells where the LTR5Hs has been homozygously or heterozygously deleted). Bottom section represents a rescue experiment, where the lack of ZNF729 gene expression is compensated by the overexpression of ZNF729 from an integrated transgene (blue). f. Representative bright field images of blastoids or defective dark spheres generated from wild type hnPSCs, ΔLTR5Hs ZNF729-/+, ΔLTR5Hs ZNF729-/-hnPSCs, or ΔLTR5Hs ZNF729-/-hnPSCs but expressing a ZNF729 transgene (rescue, ΔLTR5Hs ZNF729 + OE ZNF729). g. Quantification of the blastoid formation efficiency from wild type, ΔLTR5Hs ZNF729-/+, ΔLTR5Hs ZNF729-/-hnPSCs, or ΔLTR5Hs ZNF729 + OE ZNF729. For each condition three independent clonal cell lines were used and blastoid formation potential was measured in three independent biological replicates. h. Representative ΔLTR5Hs ZNF729-/-dark sphere immunostained with the apoptotic marker cleaved-CASP3 (red) and DAPI (blue). i. Quantification of cleaved-CASP3 immunostaining described in h (n=5). j. Representative ΔLTR5Hs ZNF729-/-dark sphere immunostained with NANOG (epiblast marker) and GATA3 (trophectoderm marker), n=3.

The remaining 107 genes had a clear ortholog in the mouse, and we analyzed their expression in the preimplantation epiblast. For simplicity, we classified the mouse orthologous genes as “expressed” or “not expressed” (Methods), with a caveat that this analysis may pass over more subtle, quantitative differences in expression between species, which can nonetheless be functionally important. We then identified genes expressed in the epiblast of both human and mice or only in humans (despite having an ortholog in mouse) (Figure 4A). We observed that among the conserved genes, 64 were expressed in humans but not in mice, whereas expression of the remaining 43 was conserved between the human and mouse epiblast. The same analysis performed using the transcriptomic data from marmosets^61^ found that, as expected, humans and marmosets share a larger proportion of conserved genes than humans and mice (only 29 genes are not conserved between marmoset and human, whereas 37 are not conserved between humans and mouse). Among the conserved genes, 53 genes are expressed only in the human epiblast and not in marmoset’s as compared to 63 in mice. Finally, marmosets and humans share a larger proportion of expressed genes in the epiblast (62 in marmosets versus 44 in mice) (Extended Data Figure 6A). In summary, our analysis shows that the cis-regulatory activity of LTR5Hs has substantially contributed to the species-specific diversification of the epiblast transcriptome, with a hundred of LTR5Hs-dependent genes being expressed in humans but not in mice and 82 being expressed in humans and not in marmosets.

### A human-specific LTR5Hs element near a primate*-*specific gene *ZNF729* is essential for blastoid formation

Considering that we established that genes with conserved expression between humans and mice are dependent on LTR5Hs for expression in humans, we speculated that they may have an ancient role in the mammalian epiblast but that the insertion of the strong LTR5Hs enhancer may have led to a relaxation of the evolutionary constraint on the ancestral cis-regulatory elements (enhancer turnover) resulting in ‘transposon addiction’^63,64^. Accordingly, we hypothesized that the observed impact of LTR5Hs on development is more likely to arise from this ‘transposon addiction’ than from evolutionarily young LTR5Hs elements regulating recently emerged genes. However, contrary to this expectation, we noted that in hnPSCs, the only LTR5Hs deletion with observable phenotype (slow growth) was the one at the *ZNF729* locus. Indeed, growth curve analyses confirmed that ΔLTR5Hs *ZNF729*^-/-^ hnPSCs exhibited a much longer doubling time of 34 h compared to 19 h for wild type cells (Figure 4B). This was surprising, considering that the LTR5Hs insertion at the *ZNF729* locus is unique to humans and not present in any other species, and that *ZNF729* gene is also evolutionary young (Zoonomia project’s Cactus genomic alignments^65,66^; Figure 4C). Although due to their repetitive nature, frequent recombination, and fast evolution, the precise evolutionary age of genes encoding KRAB zinc finger proteins (KZFPs) is challenging to determine, previous genomic analyses suggest that *ZNF729* gene emerged in the Old World anthropoids (*Catarrhini*) lineage^62,67^.

In humans, *ZNF729* expression in hnPSC and early embryos is robust and correlates with HERVK activity (Figure 4D and Extended Data Figure 6B). Interestingly, however, although the *ZNF729* gene is present in chimpanzee, our reanalysis of recently published data revealed that it is poorly expressed in chimpanzee naïve PSCs (to date, no early embryo data are available)^68^. Moreover, the *ZNF729* transcript is not present in the transcript model of macaques (Ensembl version Mmul_10) and unguided transcriptome assembly (see Methods) uncovered no evidence for *ZNF729* expression in the macaque naïve PSCs^69^. These observations suggest that expression of *ZNF729* during preimplantation development may be unique to humans and associated with the insertion of the strong LTR5Hs enhancer directly upstream from the gene. To investigate if this element may function as an alternative promoter for *ZNF729* rather than an enhancer, we examined chromatin marks at the locus in hnPSCs. This LTR5Hs element displays chromatin marks more consistent with an enhancer (high H3K27ac levels and low H3K4me3 levels) whereas the bona fide promoter shows high H3K4me3 and lower H3K27ac (Extended Data Figure 6D). Furthermore, public isoform-resolved transcriptome data from human preimplantation embryos^70^ show that *ZNF729* transcript reads originate from the *ZNF729* promoter and not from the LTR5Hs insertion (Extended Data Figure 6E).

To test the impact of this LTR5Hs enhancer deletion on the blastoid-forming potential of hnPSCs, we triggered blastoid formation from clonal cell lines lacking the LTR5Hs element at the *ZNF729* either in heterozygosity (ΔLTR5Hs *ZNF729*^-/+^) or homozygosity (ΔLTR5Hs *ZNF729*^-/-^). Strikingly, blastoids failed to form in both cases, and instead remained as dark spheres with barely any sign of cavitation, and this phenotype resembled the dark spheres we observed upon *en masse* LTRHs repression (Figure 4E, 4F, and quantified in 4G). These dark spheres also expressed the apoptotic marker Cleaved-CASP3 (Figure 4H, quantified in I), the pluripotency marker NANOG and the trophectoderm marker GATA3. GATA3 was localized to the edges of the sphere, likely because those cells are more exposed to the trophectoderm differentiation cues, still the structure failed to cavitate (Figure 4J). Importantly, blastoid formation in the ΔLTR5Hs ZNF729^-/-^ hnPSCs was partially rescued by introducing a transgene encoding *ZNF729* cDNA, indicating that ZNF729 is a key mediator of the phenotype (Extended Data Figure 6F, Figure 4E, Figure 4F and quantified in Figure 4G). We note that the incomplete rescue is likely due to a suboptimal transgene expression rather than the regulation of multiple genes by the LTR5Hs insertion, because expression of other genes at the locus is not significantly affected by the deletion (Extended Data Figure 6C). Altogether, our results indicate the deletion of a single human-specific LTR5Hs insertion regulating the primate-specific gene *ZNF729* is incompatible with blastoid formation. Thus, even highly species-specific retrotransposons can contribute to developmentally essential functions.

### ZNF729 recognizes GC-rich sequences in hnPSCs

To understand why the LTR5Hs insertion at the *ZNF729* locus is essential, we turned our attention to the gene’s product. Structurally, ZNF729 consists of the classic repressor domain KRAB and an exceptionally high – the highest in the human proteome – number of zinc-finger domains (zf-C2H2), 37 (compared to an average of 13^71^, Figure 5A). Although it has been described as a naïve and formative pluripotency marker, the molecular function of ZNF729 has not been studied^56,72^. Among genes encoding KZFPs, *ZNF729* is the most downregulated in the LTR5Hs-CARGO blastoids’ epiblast (Extended Data Figure 7A). To permit acute perturbation of ZNF729 function, we endogenously tagged the protein at the C-terminus with the dTAG-inducible degron tag FBKP12^F36V^ ^73,74^ followed by two HA tags (Figure 5A). We derived homozygously tagged clonal cell lines and confirmed that upon the addition of dTAG^v^-1, ZNF729-FKBP-HA (from now on ZNF729-FH) was rapidly degraded (Figure 5B). Next, we performed ChIP-seq of ZNF729-FH using anti-HA antibodies in DMSO control conditions and dTAG^v^-1 treated hnPSCs. We identified 46,398 regions bound by ZNF729 in hnPSCs; >95% of these peaks were lost in the dTAG^v^-1 treated sample, indicating specificity (Figure 5C and 5H, Extended Data Table 4).

**Figure 5.**
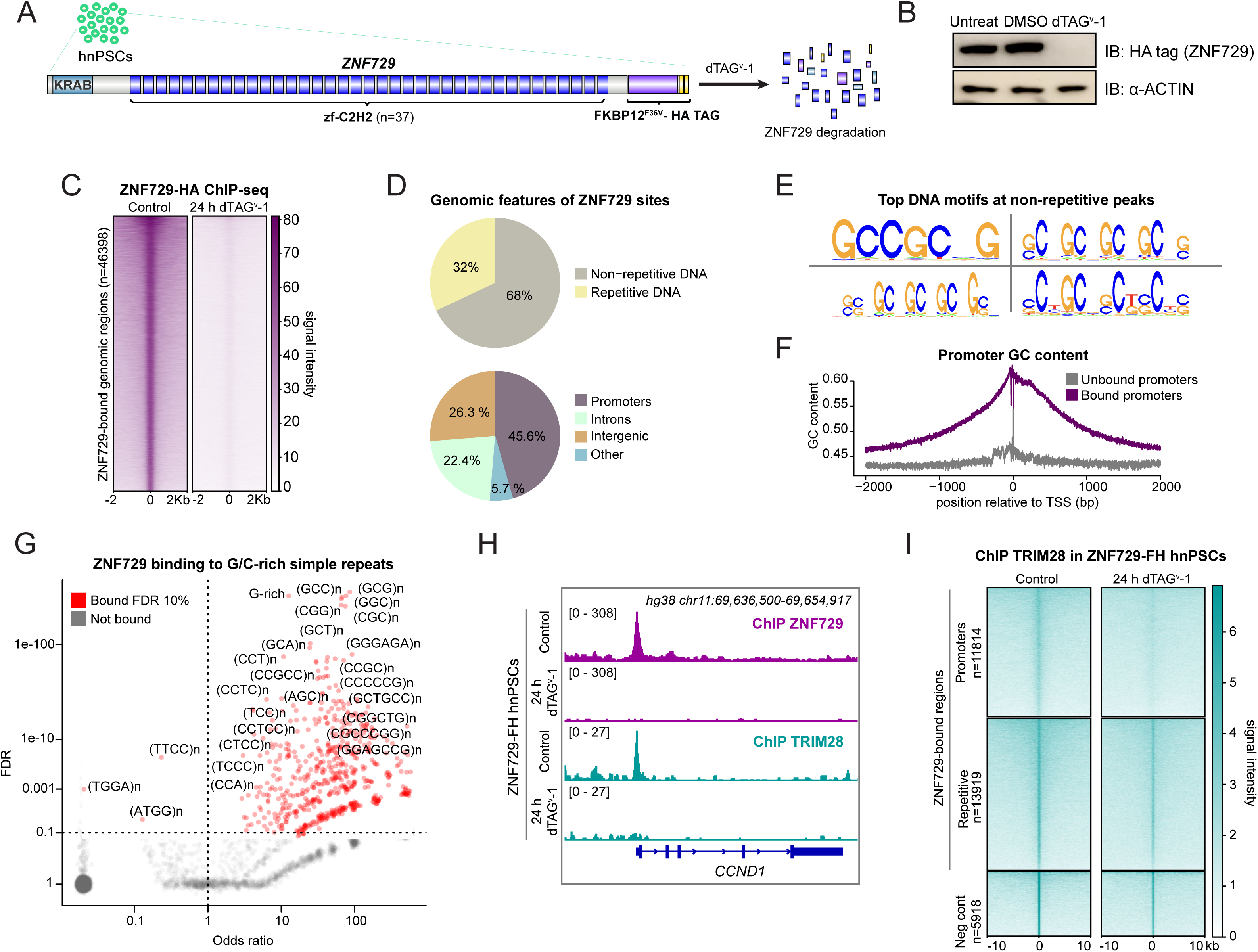
Widespread binding of ZNF729 at G/C rich sequences and promoters in hnPSC. a. Schematics depicting tagging of ZNF729, a protein containing the KRAB domain and 37 C2H2 zinc-finger domains, with an FKBPVF36V and two HA tags, in hnPSCs (ZNF729-FH hnPSCs). Upon dTAGv-1 addition, ZNF729 is degraded. Protein structure was drawn with IBS 2.0131. b. Western blot of ZNF729-FH hnPSCs untreated, treated with DMSO as control or treated with dTAGv-1. Membrane was blotted with an antibody against the HA tag and with α-ACTIN as loading control blotted on the same membrane. For gel source data, see Extended Data Figure 9. c. Heatmap displaying ZNF729-FH ChIP-seq signal over its bound genomic regions in DMSO treated (control) and 24 h dTAGv-1 treated ZNF729-FH hnPSCs (n=2). d. Pie charts displaying the genomic features of ZNF729-FH occupied regions. Top pie chart shows peak distribution between repetitive or non-repetitive DNA, bottom between promoters, intergenic or intronic DNA regions. e. Top four DNA sequence motifs obtained when performing motif discovery analysis on the ZNF729-FH ChIP-seq non-repetitive peaks using SeqPos76. f. Plot representing GC content at ZNF729-FH bound (purple line) or unbound (grey line) promoters. g. Scatter plot displays odds ratio of ZNF729-FH binding to simple repeats that contain either G/C or both. Significantly bound (FDR 10%) are depicted in red, not bound in grey. h. IGV genome browser capture of ZNF729-FH ChIP-seq signal (top two tracks, purple) or TRIM28/KAP1 ChIP-seq signal (bottom two tracks, turquoise) in DMSO treated (control) and 24 h dTAGv-1 treated ZNF729-FH. hg38 coordinates chr11:69636500-69654917. i. Heatmap displaying TRIM28 / KAP1 ChIP-seq signal over ZNF729 bound non-repetitive promoters (top), repetitive regions (middle) or regions bound by TRIM28 that do not overlap with ZNF729 (bottom). ChIP-seq signals from control-treated (left panels) or dTAGv-1 treated (right panels) ZNF729-FH hnPSCs are shown, (n=2).

KZFPs are known to bind and repress TEs via recruitment of KAP1/TRIM28 and epigenetic repressors^75^. To investigate if ZNF729 binds TEs in hnPSCs, we classified peaks based on their location. Different to most of the studied KZFPs, ZNF729 binds mostly non-repetitive DNA (68% of peaks, compared to 32% at repetitive DNA), with almost 46% of the non-repetitive peaks overlapping gene promoters (Figure 5D). At the repetitive DNA, ZNF729 binds to many TE families, and among the most enriched are young TEs, such as SVA_D/F/B, LTR12C, HERVH, L1HS and even LTR5Hs itself (Figure Extended Data Figure 7B and 7C for examples of binding in purple). Next, we performed motif discovery analysis at the non-repetitive peaks using SeqPos (Extended Data Table 5)^76^. In agreement with the KZFP DNA motif prediction still being a complex and unsolved biological problem^77–80^, the motif patterns discovered by different tools varied. Nonetheless, regardless of the specific tools used, the recovered motifs corresponded to G/C-rich sequences with diverse spacing configurations and of different length (example of discovered motifs are shown in Figure 5E). Given that mammalian promoters are enriched in the G/C-rich sequences, we wondered if the widespread binding of ZNF729 at promoters can be explained at their G/C-content. Indeed, ZNF729-bound promoters had higher G/C-content than the unbound promoters (Figure 5F). We considered that the discovery of G/C-rich motifs could be a consequence, rather than a cause of ZNF729 binding at the promoters. However, we also observed high enrichment of ZNF729 binding at G/C-rich simple repeats even in the absence of the association with promoters (FDR 10%, Figure 5G), suggesting an inherent propensity of ZNF729 to recognize G/C-rich sequences.

Given the presence of an intact KRAB domain in ZNF729, we next tested if the canonical KZFP partner KAP1/TRIM28 colocalizes with ZNF729 at promoters or elsewhere. ChIP-seq of TRIM28 in control ZNF729-FH hnPSCs confirmed colocalization of both proteins genome-wide at both repetitive and non-repetitive targets (Figure 5H and 5I). Noticeably, repetitive regions exhibited higher levels of TRIM28, consistent with multiple KZFPs likely binding those regions (Figure 5I, and Extended Data Figure 7C). Degradation of ZNF729 upon dTAGv-1 addition for 24 h broadly affected TRIM28 binding to the genome, diminishing the number of total called peaks from 17460 to 3160. The impact of ZNF729 loss was less severe at repetitive regions, again consistent with potential redundancy with other KZFPs binding at these elements. In contrast, TRIM28 binding was strongly diminished at promoters (Figure 5I). As expected, TRIM28 regions not overlapping with ZNF729 peaks were unaffected (Figure 5I). Together, these results indicate that ZNF729 recognizes G/C-rich sequences, including thousands of promoters where it recruits TRIM28.

### ZNF729 is a transcriptional regulator of basic cellular functions in hnPSCs

The association between KZFPs and TRIM28 frequently drives transcriptional silencing^75^. To investigate if ZNF729 regulates gene expression in hnPSCs, we treated the ZNF729-FH cells with dTAGv-1 for 3 h or 24 h and with DMSO as control, and performed bulk-RNA-seq using four independent endogenously tagged clonal cell lines. We performed differential gene expression analysis using DESeq2^52^ and identified differentially expressed genes at 3 h and 24 h representing acute and longer-term changes, respectively. Surprisingly, at 3 h we detected mostly gene downregulation (specifically, 270 downregulated and 6 upregulated genes) (Figure 6A), suggesting that ZNF729 may act as a transcriptional activator. Even at 24 h, there was still a larger proportion of downregulated genes (1125) than upregulated genes (559) (Figure 6B). We also quantified if TE expression was affected by ZNF729 depletion by using the software TEtranscripts combined with DESeq2^52,81^. At an FDR 5%, we did not call any statistically significant families affected (Extended Data Figure 8A), consistent with the small decrease in TRIM28 recruitment to TEs upon ZNF729 depletion (Figure 5I). This analysis searches for differences at the level of TE families, so we cannot exclude the possibility that within a family, specific individual insertions could still be impacted by ZNF729 depletion. Nonetheless, we conclude that the major impact of ZNF729 on transcriptome is at non-repetitive genes.

**Figure 6.**
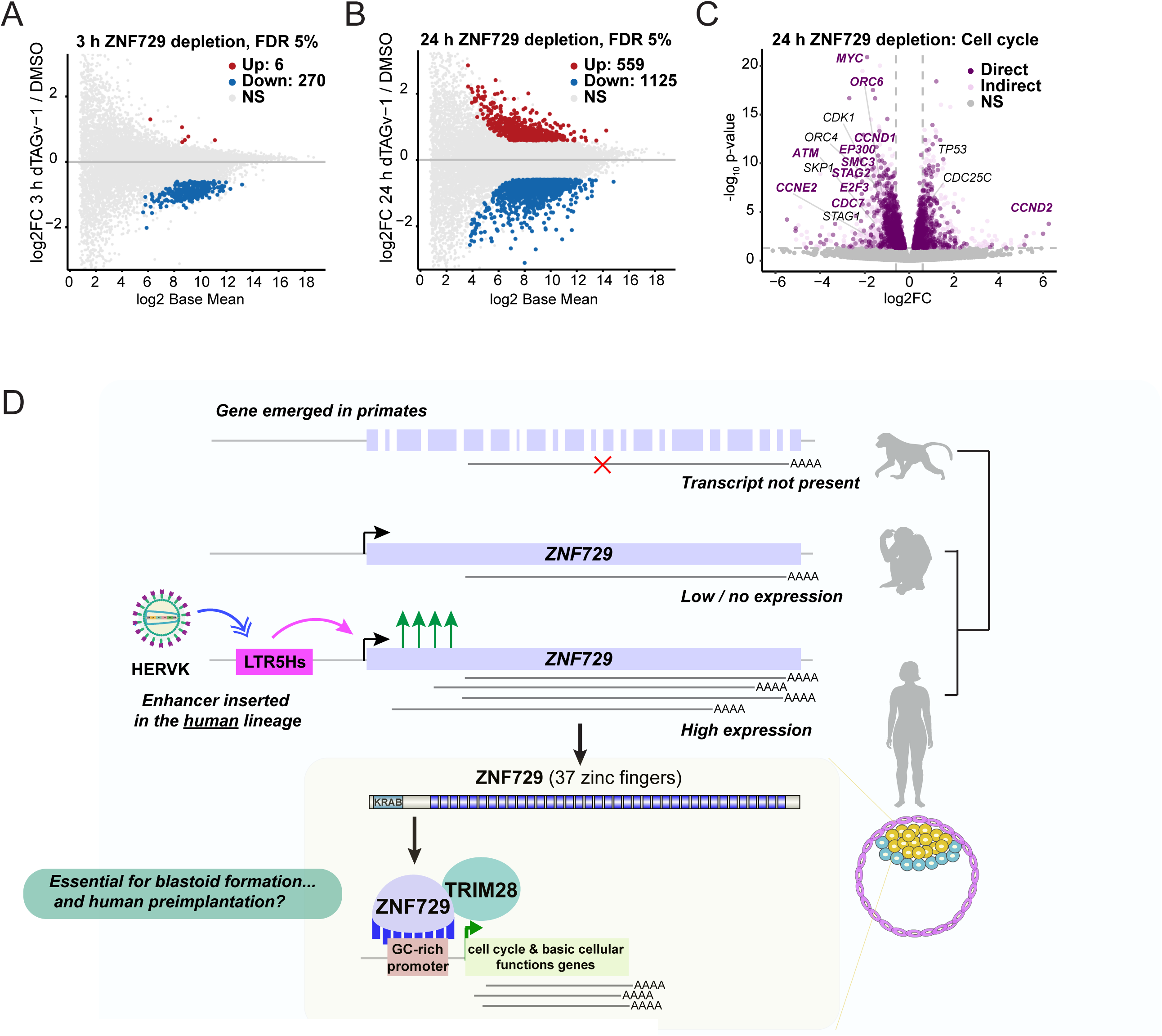
ZNF729 is a transcriptional regulator of basic cellular functions in hnPSCs. a. MA plot representing bulk RNA-seq results of ZNF729-FH hnPSCs treated with DMSO (control) or with dTAGv-1 for 3 h (n=4). Y-axis shows gene expression fold changes in 3 h dTAGv-1 treated vs DMSO treated ZNF729-FH hnPSCs. FDR 5%. b. MA plot representing bulk RNA-seq results of ZNF729-FH hnPSCs treated with DMSO (control) or with dTAGv-1 for 24 h (n=4). Y-axis shows gene expression fold changes in 24 h dTAGv-1 treated vs DMSO treated ZNF729-FH hnPSCs. c. Volcano plot of gene expression changes in bulk RNA-seq analyses (n=4) from ZNF729-FH hnPSCs treated with DMSO (control) or with dTAGv-1 for 24 h (n=4), colored by the presence (dark purple) or absence (light pink) of ZNF729-FH binding at gene promoter. Labeled dots represent genes involved in cell cycle according to GSEA curated gene sets. d. Model for human-specific gain of function of ZNF729-FH in hnPSCs, facilitated by the insertion of HERVK LTR5Hs.

We next asked if the transcriptomic changes occur at promoters directly bound by ZNF729. We defined directly regulated genes as those that were significantly misregulated upon ZNF729 loss and bound by ZNF729 within −1kb/+200 bp from the TSS, to avoid potentially confounding our results with ZNF729 bound to nearby TEs or the gene’s first intron. With these criteria, 36% of the downregulated and 35% of the upregulated gene promoters were directly bound by ZNF729, indicating that ZNF729 can act as a transcriptional activator or repressor, depending on the context (Extended Data Figure 8B and 8C). Analysis of the TRIM28 occupancy at the ZNF729-activated and ZNF729-repressed promoters upon ZNF729 depletion shows a clear reduction of TRIM28 signal in both, albeit the shape of the profiles differ between the activated and repressed promoters, suggesting that different protein complexes and configurations may be driving these activities (Extended Data Figure 8D). These results demonstrate that ZNF729 is a novel transcriptional regulator in hnPSCs.

Promoters of genes involved in basic cellular functions (sometimes referred to as ‘housekeeping’ promoters) are typically GC-rich and highly and broadly active across cell types^82,83^. Intriguingly, gene promoters bound by ZNF729 are not only G/C-rich, but also display median levels of expression five times higher than the unbound promoters (Extended Data Figure 8B). Moreover, ontology analysis of genes affected by ZNF729 loss at 3 h, when changes are mostly direct, are associated with basic cellular processes like cell division, regulation of GTPase activity, regulation of RNA metabolic process and others (Extended Data Table 6). The cell division category led us to reason that the slow growth phenotype of ΔLTR5Hs *ZNF729*^-/-^ hnPSCs could be explained by the cell cycle genes being affected upon ZNF729 depletion. Indeed, systematic analysis of cell cycle related genes using a curated gene set from GSEA^41^ revealed that classic cell cycle regulators such as *MYC*, *CDK1*, *CCND1* among others are directly bound and activated by ZNF729 (Figure 6C and Figure 5H for a browser capture showing the *CCND1* gene). These results suggests that ZNF729, with its key role in regulating basic cellular processes in hnPSCs, may be a major driver of phenotypes observed upon LTR5Hs repression. Consistent with this notion, PCA revealed similarities between transcriptomes of ZNF729-depleted hnPSCs and those of dark spheres formed from the high repression LTR5HS-CARGO cells (Extended Data Figure 8E). Altogether, our results demonstrate how the emergence of a new gene during primate evolution, the subsequent capture of a retrovirus-derived enhancer in the human lineage that allowed for its expression, and the acquisition of target specificity of ZNF729 towards GC-rich sequences (extremely abundant in the mammalian promoters), influenced essential cellular programs during early human development (Figure 6D).

## Discussion

Human embryo research remains crucial for establishing the ground-truth reference, however, functionally malleable 3D embryo models overcome major ethical and practical challenges associated with the human embryo work and offer an accelerated path for systematic discovery of mechanisms underlying human embryogenesis^84^.

By repressing HERVK LTR5Hs *en masse*, we demonstrate that the activity of the hominoid-specific retrotransposon HERVK LTR5Hs is required for proper blastoid formation and lineage identity. LTR5Hs elements function as enhancers for many genes, some of them established markers of human epiblast/naive pluripotency such as *SUSD2* and *ZNF729* ^31,42,56^. Consequently, LTR5Hs repression severely affects the blastoids’ epiblast transcriptome, leading to downregulation of many genes in cis and to an upregulation of genes associated with trophectoderm and placenta development. We postulate that the latter effect is the by-product of the combined loss of fidelity of the naive epiblast state (i.e. widespread changes in gene expression in the blastoids’ epiblast) and the presence of signals promoting trophectoderm differentiation during blastoid formation. LTR5Hs repression also leads to a diminished allocation of cells into the hypoblast and polar (but not mural) trophectoderm lineages. Given that in embryos specification of the hypoblast and polar trophectoderm depends on the epiblast^26,38^, we speculate that these effects may be an indirect result of the blastoids’ epiblast being defective and displaying impaired maturation (Figure 2F). Nonetheless, HERVK LTR5Hs is active in the hypoblast too, therefore we cannot rule out that hypoblast-autonomous LTR5Hs effects contribute to the exhaustion of this population.

The impact of HERVK LTR5Hs for blastoid formation likely results from convergence of multiple mechanisms. One, explored here, is its cis-regulatory activity: many genes expressed in the epiblast are regulated by LTR5Hs in cis, often at long ranges, and we demonstrate that at least one LTR5Hs element – that at the *ZNF729* locus – is essential for blastoid formation through its cis-regulatory function. Some of the other LTR5Hs-dependent genes may confer essential functions, whereas others may have no impact on early development. Beyond the cis-regulatory activity of the LTR5Hs, HERVK LTR5Hs insertions are transcribed, still retain different levels of coding capacity for retroviral proteins^85^ and these proteins can assemble viral-like particles in human blastocysts^15,86^. These RNAs and retroviral products might also have regulatory roles in the preimplantation embryo ^87–95^. However, our observations that overexpression of HERVK retroviral proteins does not rescue the failed blastoid formation suggest that at least for this phenotype, the proteins themselves do not play a major role (Extended Data Figure 2E). Nonetheless, these proteins may play other roles in the embryo and our study represents only the first step towards unpicking the complexity of HERVK contributions to human preimplantation development.

Comparing transcriptomes of human, marmosets and mice, we highlight a substantial contribution of HERVK LTR5Hs to the hominoid-specific diversification of the epiblast transcriptome. One of the divergently expressed genes is the primate-specific *ZNF729* gene. We reveal a requirement for the human-specific LTR5Hs insertion controlling the *ZNF729* expression in hnPSCs proliferation and blastoid formation potential. Mechanistically, ZNF729 confers this essentiality by binding and regulating hundreds of crucial genes due to its affinity for G/C-rich sequences, which are extremely abundant in mammalian promoters^82^. ZNF729 contains an intact KRAB domain, capable of mediating transcriptional repression in a reporter screen of human transcriptional effectors^96^. In agreement, we show that ZNF729 is the major transcription factor responsible for recruiting TRIM28, the KRAB-domain binding corepressor, to promoters in hnPSCs. Surprisingly, however, a substantial fraction of the regulated promoters are activated, rather than repressed by ZNF729. While the exact mechanism underlying this activation remains to be established in future studies, we note that in addition to its co-repressor function, TRIM28 has also been implicated in gene activation through mediating RNA polymerase II (RNAPII) pause-release^97–99^. In that context, it is also interesting to note that RNAPII preferentially pauses at GC-rich sequences^100,101^. Alternatively, ZNF729 activator function may be mediated by a heretofore unknown TRIM28-independent mechanism.

Regardless of specifics, a fundamental question remains: why is an evolutionary young gene, controlled by an even younger, human-specific enhancer, important for regulation of ancient cellular programs, such as cell proliferation? We speculate that during evolution, the expansion of the zinc finger array and mutations at amino acid residues contacting the DNA, ultimately led to high specificity and affinity of ZNF729 for G/C-rich sequences. Given that such sequences are extremely enriched at promoters, especially those associated with the housekeeping functions, this G/C specificity targeted ZNF729 to such promoters, in turn boosting their activity via pause-release or other mechanism and providing a competitive proliferative advantage to cells expressing ZNF729. But such regulation could have also led to a relaxation of the constraint on the ancient mechanisms controlling expression of proliferation genes, with ZNF729 ultimately supplanting some of these mechanisms and becoming essential. Beyond ZNF729, a recent preprint highlights that other primate-specific KZFPs such as ZNF519 may also regulate cell cycle progression^102^. Thus, while ZNF729 expression appears to be limited to the early embryo and testis, the principles uncovered in our study may be relevant in other biological contexts. Moreover, recent work showed that another primate-specific Zinc-finger protein, ZNF808, is essential for proper pancreas development in humans^103^. These observations suggest that evolutionary remodeling of gene regulatory networks can result not only in species-specific innovation, but also create new dependencies and bestow essentiality on recently emerged cis-regulatory elements and genes.

## Methods

### Ethics

The use of blood-derived induced naive pluripotent stem cells for the experiments described in this manuscript was approved by the Stanford Stem Cell Research Oversight committee (SCRO Protocol number 900).

### Cell culture

NANOG and KLF2-induced hnPSCs generated from peripheral blood cells by overexpressing^34,35^ were grown on irradiated Cf1 mouse embryonic fibroblast feeder layers (MEFs) (Fisher Scientific, A34181). Prior to experiments entailing next-generation sequencing, hnPSCs were plated without MEFs (feeder-free conditions) using Geltrex (Gibco, A1413301) as a matrix. hnPSCs were grown in PXGL^42^. This medium consists of N2B27 as base, which is made by mixing 1:1 DMEM/F-12 (Sigma, D8437) and Neurobasal (Thermo, 21103049) with the following added supplements: 2mM L-glutamine (Thermo, 25030024), 100 uM 2-Mercaptoethanol (Sigma, M3148), N2 and B27 supplements (Gibco, 17502048 and 7504044), and 1x Antibiotic-Antimycotic solution (Sigma-Aldrich A5955-100ML). To make PXGL, we freshly supplemented the following chemicals: 1 uM PD0325901 (Selleckchem, S1036), 2 uM XAV939 (Cell Guidance Systems, SM38-200), 2uM Go 6983 (Bio-Techne, 2285), 10ng/mL recombinant human LIF (Preprotech 300-05), and 1 ug/mL of doxycycline to sustain NANOG and KLF2 transgenes expression in hnPSCs. Doxycycline was eliminated from the media for the nontarg-CARGO or LTR5Hs-CARGO induction experiments. Cells were passaged using Tryple Express (Fisher Scientific 12-605-010) every 3-4 days or whenever colonies were too confluent. The cell incubator was kept at 37 degrees, humidified, at 7% CO2 and 5% O2 (hypoxia). All cell lines were tested monthly for Mycoplasma. For nontarg-CARGO and LTR5Hs-CARGO induction we used 2x water soluble cumate (System Biosciences QM150A-1).

### Derivation of nontarg- and LTR5Hs-CARGO hnPSCs and rescue with HERVK ORFs

hnPSCs were nucleofected using a Lonza 4D-Nucleofector using the P3 Primary X Kit-S (Lonza, V4XP-3032), and the DN100 program. Per nucleofection we used 400,000 cells without MEF depletion. To generate the KRAB-dCas9 hnPSCs, 0.8 ug of a piggyBac construct containing KRAB-dCas9 under a cumate-inducible promoter and a puromycin selection cassette were co-nucleofected with 0.2 ug of the super piggyBac transposase (System Biosciences, PB210PA-1). Clones containing the integration were selected with puromycin (0.5ug/mL) for three passages. KRAB-dCas9 hnPSCs cells were later nucleofected with 0.8 ug of the piggyBac constructs containing the nontarg-CARGO (Addgene #191319^17^) and the LTR5Hs-CARGO (Addgene #191316^17^) and a neomycin selection cassette and 0.2 ug of the super piggyBac transposase. Cells were then selected with 200 ug/ml of G418 for 10 days. 2000 cells were subsequently plated in a 10cm^2^ plate containing MEFs and fed every day. On day 8-9 sparse colonies are visible and were picked and expanded for the experiments. Cells were treated with puromycin and G418 every few passages to sustain proper KRAB-dCas9 and CARGO array expression, as we noticed these transgenes get silenced over the passages in hnPSCs. For the ‘orthogonal repression of LTR5Hs’ experiments, a distinct array of guide RNAs targeting LTR5Hs was designed and cloned into piggyBac using CARGO^39^ (gRNA sequences in Extended Data Table 7). The LTR5Hs-Ortho-CARGO hnPSCs were generated as described above, with the only difference that this time the KRAB-dCas9 transgene was under a cumate-inducible Ef1a promoter to ensure high repression at the population level. Analysis of the role of the HERVK proteins in the dark spheres phenotype (‘rescue with HERVK ORFs’ experiment) was performed by selecting three high repression LTR5Hs-CARGO clones that were previously demonstrated to give rise to dark spheres, and integrating into them a piggyBac transgene encoding a tagBFP and the proteins gag, pro, and pol^40^ under a constitutive Ef1a promoter to ensure robust expression. High repression LTR5Hs-CARGO hnPSCs positive for tagBFP were isolated and utilized for blastoid formation under cumate treatment to induce LTR5Hs-repression.

### Genetic deletion of LTR5Hs elements and ZNF729-HA overexpression

Selected LTR5Hs elements were deleted from the genome using pairs of gRNAs designed using Benchling [Biology Software]. (2023), (Extended Data Table 7). crRNAs were purchased from IDT with the XT modification for stability. 400,000 cells were nucleofected with a ribonucleoprotein complex containing 1.65 ug of HiFi Cas9 Nuclease V3(IDT, 1081059) and 0.85 ul of a 1:1 ratio of 100uM annealed tracRNA and crRNA. Cells were passaged once and then 2000 cells were plated on a 10cm^2^ plate with MEFs for colony picking. Clones were genotyped using PCR and Sanger sequencing, and heterozygous and homozygous clones were kept for experiments. For the rescue experiment described in Figure 4E, 400,000 ΔLTR5Hs *ZNF729*^-/-^ hnPSCs were nucleofected with a piggyBac plasmid subcloned from a pcDNA3 vector, containing ZNF729-HA cDNA (purchased from Genscript) and a puromycin selection cassette. Super piggyBac transposase was co-nucleofected. Cells were selected with 0.5ug/mL puromycin for 10 days and ZNF729-HA expression was tested by western blot.

### Derivation of ZNF729-FKBP^F36V^-HA hnPSCs and dTAGv-1 treatments

To endogenously tag *ZNF729*, we performed homology-directed repair at the locus with a donor DNA providing the FKBP^F36V^ and HA tags. To this end, we drew upon a previously published method^104^ based on the combination of Cas9 ribonucleoproteins and delivery of the donor template by AAV6 viral vectors. To generate the AAV viral particles, 2 x 15cm^2^ dishes of 293FT cells at 60% confluency were transfected. The day of transfection, the 293FT cells “complete cell media” (DMEM/High Glucose Medium, Cytiva SH30243.FS; 10% FBS, GeminiBio 100-106; 1X non-essential amino acids,Gibco 1114-0050; 1X GlutaMAX, Gibco 4109-0036; 1X Antibiotic-Antimycotic, Gibco 1524-0062) was refreshed 6 hours before transfecting. Transfection was carried out using 120 ug polyethylenimine (PEI) per 15 cm^2^ plate, 22ug of pDGM6 (Addgene 110660^105^) and 6ug of AAV template (cloned in the pAAV-GFP backbone, Addgene 32395^106^). After 24 h, the media was changed to “slow growth media” (same as complete media, but with 2% FBS instead of 10%) and upon further 48 h of culture, the AAV viral particles were purified using one reaction of the AAVpro kit (Takara Bio 6675) and stored at −80 degrees. The crRNA (Extended Data Table 7) to target the *ZNF729* C-terminal region was purchased from IDT with the XT modification for stability. 400,000 wild type hnPSCs were nucleofected with the ribonucleoprotein complex containing 1.65 ug of HiFi Cas9 Nuclease V3 (IDT, 1081059) and 0.85 ul of a 1:1 ratio of 100uM annealed tracRNA and crRNA. Cells were seeded in a plate containing MEFs, PXGL, the ROCK inhibitor Y-27632, and the AAV viral particles containing the donor template. Media was changed after 24 h. Cells were passaged once and then 2000 cells were plated on a 10cm^2^ plate with MEFs for colony picking. Correct editing was analyzed by PCR, Sanger sequencing, and western blot. ZNF729 depletion was obtained upon addition of 500 nM of dTAG^v^-1 for the indicated times (Tocris 6914).

### Blastoid formation

To generate blastoids the protocol described in Kagawa et al.^26,33^ was followed with minor changes. hnPSCs were grown on MEFs and dissociated the day of the experiment into single cells. MEFs were depleted by culturing the dissociated cells in PXGL over a gelatin matrix (Sigma-Aldrich G1393) for 1 h. We used 24-wells Aggrewell 400 (StemCell Technologies 34415) plates as vessels. Upon multiple tests, we determined that starting from 76 cells per intended blastoid was optimal, so 91,200 hnPSCs were plated per well of the microwell plate (76 multiplied by 1200 microwells). On the day of plating, cells were cultured in N2B27 base medium containing 10 uM Y-27632 (StemCell Technologies, 72304). After 20-24 hours, medium was changed to PALLY medium (N2B27 base medium supplemented with PD0325901 (1 uM), A83-01 (1 uM, MedChemExpress, HY-10432), 1-Oleoyl lysophosphatidic acid sodium salt (LPA) (500nM, Tocris, 3854), hLIF (10 ng/mL), and Y-27632 (10 uM). PALLY medium was refreshed the next day. 72 h after plating, medium was replaced with medium containing 500nM of LPA. At 96 h, structures were collected and analyzed as needed.

### hnPSCs differentiation towards the trophectoderm lineage

Trophectoderm monolayer differentiation was completed as described previously^50,107^. Briefly, hnPSCs were washed with PBS and then incubated with TrypLE Express for 10 min at 37 degrees. Dissociated cells were washed in DMEM/F-12 (Thermo Fisher Scientific, #11-330-057) with 0.1% Bovine Albumin Fraction V (ThermoFisher Scientific #15260037) and resuspended in nTE-1 media (N2B27 media supplemented with 2 uM PD325901, 2 uM A83-01, and 10 ng/mL BMP4 (R&D Systems 314-BP-010)). Cells were counted and seeded to plates coated with 0.15 ug/cm^2^ laminin511-E8 (Amsbio AMS.892 021) at a density of 2×10^4^ cells per cm^2^. 24 h after plating, media was changed to nTE-2 media (N2B27 media supplemented with 2 uM PD325901, 2 uM A83-01, and 1 ug/ml JAK inhibitor I (StemCell Technologies 74022). 48 h after plating, media was again changed to fresh nTE-2 media. To repress LTR5Hs elements during the differentiation, media was supplemented with 2x water soluble cumate. Differentiations took place under hypoxic conditions.

### hnPSCs differentiation towards the hypoblast lineage

Hypoblast monolayer differentiation from hnPSCs was completed as described previously^51^. Briefly, hnPSCs were washed with PBS and then incubated with TrypLE Express for 10 min at 37 degrees. Dissociated cells were washed in DMEM/F-12 with 0.1% Bovine Albumin Fraction V and resuspended in a six-factor “6F media” (N2B27 media supplemented with 25 ng/mL FGF4 (PeproTech 100-31; stabilized with 1 µg/mL heparin sodium), 10 ng/ml recombinant human BMP4, 10 ng/ml recombinant human PDGF-AA (Peprotech, 100-13A), 1 uM XAV939 (Cell Guidance Systems SM38-10), 3 uM A83-01 (MedChem Express HY-10432) and 0.1 uM retinoic acid (Sigma-Aldrich R2625). Cells were counted and seeded to plates coated with 0.15 µg/cm^2^ laminin511-E8 at a density of 5×10^4^ cells per cm^2^. 24 h after plating, the medium was replaced with fresh 6F media. 48 h after plating, the medium was changed to a seven-factor “7F media”, which includes the same factors used in the 6F media, with the addition of 10 ng/mL recombinant human IL-6 (PeproTech 200-06). To repress LTR5Hs elements during the differentiation, media was supplemented with 2x water soluble cumate. Differentiations took place under hypoxic conditions and flow cytometry measures were taken on day 3.

### Flow cytometry

After 3 days of trophectoderm or hypoblast differentiation, 200,000 cells were used for staining. Cells were pelleted and resuspended in 100 uL of N2B27 supplemented with 10 uM Y-27632 and either a 1:100 dilution of TACTSD2-BV421 for the trophectoderm differentiations (BD Biosciences 563243) or 1:200 dilution of ANPEP-BV421 for the hypoblast differentiations (BioLegend 301716). Cells were incubated on ice in the dark for 1 h and then washed twice with N2B27 supplemented with 10 uM Y-27632. Flow cytometry was performed on the SONY MA900 cell sorter and data were analyzed using FlowJo v.10.10.0.

### RNA extraction and RT-qPCR

RNA extraction was performed using Trizol (Life Technologies, 15596018) directly on dissociated hnPSCs carrying the indicated perturbation or, in the case of blastoids and dark spheres, prior to RNA extraction, the structures were dissociated in a 1:1 mixture of trypsin-EDTA 0.5% (Fisher Scientific, 15-400-054) and Accutase (StemCell Technologies, 07920) for 5 min, diluted in N2B27 and spun down. Extraction was followed by RNA purification using a Direct-zol RNA-prep kit (Zymo Research, R2052) with DNAse treatment. 1 ug of RNA was retrotranscribed into cDNA using a SensiFAST cDNA synthesis kit (Bioline, BIO-65053), cDNA was diluted 1:4 with molecular grade water and 2 ul of this dilution were used for qPCR with primers for each amplicon (Extended Data Table 7). qPCR was performed in a LightCycler 480 Instrument (II) using a SensiFAST SYBR (Bioline, BIO-98020). For experiments using Taqman probes, qPCR Primetime probes were purchased from IDT (sequences in Extended Data Table 7) and were combined for qPCR with the LightCycler 480 Probes Master mix (Roche, 04707494001).

### CUT&RUN

Protocol was performed according to Meers et al.^108^ and using the CUTANA reagents from EpiCypher (Concanavalin A conjugated paramagnetic beads, 21-1401; pAG-Tn5, 15-1017; E.coli spike-in DNA, 18-1401). We used 500,000 nontarg-CARGO or LTR5Hs-CARGO hnPSCs per condition and permeabilized using 0.005% of digitonin, 0.5 ug of H3K9me3 primary antibody were used per sample (Extended Data Table 7). DNA was extracted using phenol-chloroform and library preparation was performed using the NEBNext Ultra II Library Prep kit (New England Biolabs, E7645S). Libraries were sequenced paired-end 150 cycles in a Novaseq 6000 Illumina sequencer.

### ChIP-seq, ChIP-seq libraries construction, and sequencing

Cells were grown on feeder-free conditions using Geltrex to minimize MEF contamination in the sequencing. One 10 cm^2^ (∼6×10^6^ hnPSCs) was used per chromatin immunoprecipitation. Cells were crosslinked in PBS containing 1% methanol-free formaldehyde (Pierce, 28908) for 10 min. Fixation was quenched during 10 min by adding a final concentration of 0.1M of glycine. Upon harvesting, cells were resuspended in buffer 1 (50 mM HEPES-KOH pH 7.5, 140 mM NaCl, 1 mM EDTA, 10% glycerol, 0.5% NP40, 0.25% Triton X-100) and incubated for 10 min, rotating at 4 degrees, prior to centrifugation at 1350xg for 5 min at 4 degrees. The pellet was lysed in buffer 2 (10 mM Tris pH 8, 200 mM NaCl, 1 mM EDTA, 0.5 mM EGTA), incubated for 10 min at 4 degrees and once again centrifugated at 1350xg for 5 minutes. Then, the pellet was lysed in buffer 3 (10 mM Tris pH 8, 100 mM NaCl, 1 mM EDTA, 0.5 mM EGTA, 0.1% sodium deoxycholate, 0.5% N-lauroylsarcosine), incubated for 20 min on ice and sonicated in a Bioruptor sonicator (Diagenode) until the obtention of DNA fragments of sizes ranging 400-600 bp. Chromatin was quantified, and ∼10 to 25 ug of chromatin were used for immunoprecipitation in a total of 500 ul of buffer 3 containing the antibodies indicated in Extended Data Table 7. After overnight incubation, 100 ul of magnetic protein G beads (Life Technologies, 10004D) were added to each immunoprecipitation. After 2-3 h of incubation, the immunocomplexes were washed five times with RIPA wash buffer (50 mM HEPES-KOH pH 7.5, 500 mM LiCl, 1 mM EDTA, 1% NP40, and 0.7% sodium deoxycholate) and once with TE-NaCl buffer (50 mM Tris pH 8, 10 mM EDTA, and 50 mM NaCl). To recover the DNA, the immunocomplexes were eluted in elution buffer (50 mM Tris pH 8, 10 mM EDTA, and 1% SDS) at 65 degrees for 15 min with vortexing every 5 min. The beads eluate was decrosslinked overnight at 65 degrees. After RNAse A treatment for 30 min (Thermo Fisher Scientific FEREN0531) and proteinase K treatment (Thermo Fisher, EO0492) for 2 h, the DNA was purified using a Qiagen kit (Qiagen 28106).

To prepare ChIP-seq libraries for sequencing, we utilized the NEBNext Ultra II DNA kit (NEB, E7645S) kit and Agencourt AMPure XP beads (Beckman coulter, A63881) were used for the cleanings. We started from ∼20-50 ng of ChIP or input DNA and followed manufacturer’s instructions. Paired-end sequencing (150 cycles) was performed in a Novaseq X Plus sequencer (Illumina) including 1% of PhiX.

### Bulk RNA-seq and library preparation

RNA was extracted using Trizol from nontarg-CARGO and LTR5Hs-CARGO hnPSCs treated with cumate during four days in the absence of doxycycline, from ZNF729-FH hnPSCs treated with dTAG^v^-1 for 3 and 24 h or from blastoids and dark spheres. Messenger RNA was purified using poly-T oligo-attached magnetic beads. After fragmentation, the first strand cDNA was synthesized using random hexamer primers followed by the second strand cDNA synthesis. Libraries were prepared by end repair, A-tailing, adapter ligation, size selection, amplification, and purification and they were checked with Qubit and qPCR for quantification and Bioanalyzer for size distribution detection. Quantified libraries will be pooled and sequenced on a Novaseq 6000 Illumina sequencer.

### Blastoid immunostainings

Immunostaining of blastoids was performed *‘in well’*. Media from Aggrewell was carefully aspirated (more than 90%). For fixation, 1 mL of 4% paraformaldehyde was added to the well and incubated at room temperature for 15 minutes. The paraformaldehyde was carefully aspirated and substituted for a rinse buffer composed of PBS with 3mg/mL polyvinylpyrrolidone (PVP). Blastoids were left in PBS-PVP buffer for 5 minutes to ensure they sediment before aspirating the PBS-PVP buffer. After one rinse, blastoids were permeabilized in PBS-PVP containing 0.25% of Triton X-100 for 30 minutes. Permeabilization solution was aspirated substituted with blocking buffer (0.1% BSA (Sigma-Aldrich, A9418), 0.01% Tween 20 (P1379), 2% donkey serum (Jackson Immunoresearch, 017-000-121) which was dispensed in the well with a 5 mL serological pipette to subsequently collect all the blastoids from the well and deposit them into a well of a 6-well plate containing more blocking solution. Blocking took place for at least 3 hours at 4 degrees. Blastoids were picked using standard mouth pipetting or 20 ul pipette tips and moved to primary antibodies (Extended Data Table 7) diluted in blocking solution in Nunc MicroWell MiniTrays (Fisher Scientific 12-565-154) at 4 degrees overnight. Blastoids were washed three times with blocking buffer and stained with Alexa Fluor secondary antibodies for 3 h, washed three times and imaged in blocking buffer using an 18-well microslide (Ibidi, 81826) in an Inverted Zeiss LSM 780 confocal microscope.

### PIP-seq

PIP-seq is an alternative to microfluidics-based scRNA-seq methods that captures cells via vortex and can be performed from beginning to library preparation at the experimenter’s bench. Blastoids from two wells of an Aggrewell plate per condition were collected on a 15 mL tube, were centrifuged for 2 min at 250 xg, and media was aspirated. Blastoids were then resuspended in Collagenase IV (StemCell Technologies, 07909) and incubated at 37 degrees with mild agitation for 40 min. Blastoids were centrifuged again in N2B27 medium and the pellet was resuspended in 0.5% Trypsin-EDTA (Fisher Scientific, 15-400-054) and incubated for 10 min at 37 degrees. Two further washes were performed with N2B27, and the dissociated cells were passed through a 40 um Flowmi cell strainer. Experiment only continued when viability was larger than 80%. 40,000 cells or less were counted, captured, and used for completing the PIP-seq T20 3’ Single Cell RNA Kit protocol (Fluent Biosciences, FBS-SCR-T20-4-V4.05) without changes and using 12 cycles of cDNA amplification. Libraries were prepared with the reagents in the kit and were sequenced in an Illumina Novaseq X instrument.

### Western blot

After SDS-Page electrophoresis, protein transfer is carried out on a multilayered cassette including a nitrocellulose membrane. The transfer buffer was composed of 25mM Tris-HCl, 192mM glycine, 0.05% SDS and 10% methanol. The power source was set to 125V for 90 minutes. The nitrocellulose membrane was blocked with 5% milk for 1 hour and incubated HA tag antibody (Extended Data Table 7) overnight to detect ZNF729-HA or ZNF729-FH.

### Image obtention and quantification

All bright field images were taken using the EVOS FL Imaging System. The fluorescent immunostainings were imaged using an Inverted Zeiss LSM 780 confocal microscope. To obtain blastoids ICM/TE ratios, we used the Fiji software^109^ to measure the diameter of the blastoids cavity and the ICM size by measuring the distance from the point of contact with the trophectoderm to the end of the ICM. To count number of cells expressing specific lineage markers we used a combination of software and manual counting. KLF17, GATA4, and cleaved-CASP3 positive cells were counted with Fiji’s cell counter in each stack. GATA3 positive cells were counted using the 3D Object Counter plug-in from Fiji, carefully curating the assigned positive cells with the human-eye positive detected cells and correcting when necessary (e.g. fluorescent artifacts that are not cells).

### Quantification of blastoid formation efficiency

For determining blastoid efficiencies, endpoint (96 h) blastoids were moved to a 15 mL conical tube and the total volume was measured. Next, two technical duplicates of 50 ul aliquots were dispensed into a 96-well plate and the structures were evaluated and counted, ultimately extrapolating to the total conical tube volume and to the 1200 microwells present in the Aggrewell. To consider a 3D structure a blastoid we followed previously established criteria^26^. Briefly, its morphology should resemble stage B6 of human blastocyst, with an accumulation of cells surrounded by a monolayered cyst mimicking the inner cells mass and the trophectoderm respectively. The blastoids’ inner cell mass is often outside the cyst, in such case, we still consider that structure a blastoid. Blastoids have an approximate total diameter between 150 and 250 um, and in the case of LTR5Hs-CARGO blastoids the cavity should be larger than 150um, with no upper limit. When tested by scRNA-seq or immunofluorescence, blastoids must express markers consistent with the blastocyst’s lineages. Dark spheres are structures that appear darker in bright field and are not cavitated.

### Recording of blastoid formation movies

The blastoid formation protocol was performed as indicated above, with changes. Instead of using Aggrewells (which have an opaque bottom) we utilized Elplasia 24-well plates that allow imaging from below (Corning, 4441). We note that the initial cell aggregation in these plates is not as robust, thus end-point blastoid formation is less efficient than in Aggrewells. Cell aggregates are monitored, and when there are early signs of cavitation (small ‘bubbles’ around the aggregates), the plate is moved to a Nikon Eclipse Ti-E microscope that is equipped with a system for CO_2_ and temperature control (OKOlab). Blastoids were imaged at 37 degrees and 5% of CO_2_ for 24 h.

### CUT&RUN analysis

Standard Illumina adapters were cut from the Illumina reads using Cutadapt^110^ and then aligned to a combined hg38 and E. coli genome version using Bowtie2^111^, with the -dovetail parameter on and the rest of parameters in its default behavior. This means that in case of multimapping all the valid alignments are reported. PCR duplicates were removed from the analysis. Coverage bigwig files were generated with Deeptools^112^ bamCoverage and the –scaleFactor was set to the number obtained from the normalization of fragments mapped to the human genome (hg38) and the mapped fragments to the E. coli k12 MG1655 genome. Browser captures were obtained from IGV^113^.

### ChIP-seq analysis

Standard Illumina adapters were cut from the Illumina reads using using Cutadapt^110^. Reads were aligned to the *Homo sapiens* (hg38) genome using Bowtie2^111^ in its default behavior. PCR duplicates were removed from the analysis using Samtools^114^. Coverage bigwig files were generated with Deeptools^112^ bamCoverage. Browser captures were obtained from IGV^113^. Peaks were called using MACS3^115^. Identification of ZNF729-FH-bound repetitive DNA was performed by intersecting ZNF729-FH peaks with RepeatMasker (RRID:SCR_012954)^116^ using Bedtools^117^ intersect with -f 0.3. To be considered a peak at the promoter, ZNF729-FH or TRIM28 must bind −1kb / +200 bp around the TSS.

### RNA-seq and gene ontology analysis

Illumina adapters were trimmed from reads using Skewer^118^. Transcript alignment and quantification was performed using Salmon^119^ against the human genome assembly version Gencode v47^120^. For differential gene expression analysis we used DESeq2^52^ after excluding transcripts with less than 10 reads across the tested samples. DESeq2 compared the effect of nontarg-CARGO and LTR5Hs-CARGO in the hnPSCs, the differences between blastoids and dark spheres, or the gene expression changes upon dTAG^v^-1 addition to the ZNF729-FH hnPSCs. Biological replicates were used as covariates. Analysis of chimpanzee naïve pluripotent stem cells bulk RNA-seq^68^ was performed in the same manner but using the *Pan troglodytes* panTro6 Clint_PTRv2 genome assembly. Rhesus genome Mmul_10 (RheMac10) genome reference lacks a *ZNF729* transcript model. To assess presence and expression of *ZNF729* in rhesus macaques naive state, we performed unguided transcriptome assembly from rhesus monkey (*Macaca mulatta*) naive pluripotency stem cells bulk RNA-seq data^69^ using the Trinity pipeline^121^. We constructed a blast database from the Trinity output and searched for the human *ZNF729* nucleotide sequence using blastn and tblastx algorithms^122^. Highest matches were searched against non-redundand NCBI database with blastn and blastx algorithms. None of the sequences scored *ZNF729* in reciprocal blast as a top match. When a sequence had a blast matching to *ZNF729*, such matches were below 60% identity. Differentially regulated genes (FDR 5%) were used for human gene ontology analysis using Gorilla^123^ using as a background the list of genes expressed in hnPSCs, blastoids, and dark spheres.

### TEtranscripts

The software ‘TEtranscripts’^81^ was used to find differentially regulated TEs in the ZNF729-FH dTAG^v^-1 bulk RNA-seq experiments. To this end, the RNA-seq reads were aligned using HISAT2^124^, and following the tool’s manual, we allowed 100 alignments per read (-k 100) to optimize TE quantification and differential analysis.

### PIP-seq and pseudobulk differential gene expression analysis

For each sample and replicate, reads obtained from the Novaseq X were analyzed with Fluent Bio’s proprietary software Pipseeker with default parameters and aligning against the GRCh38 transcriptome index (Gencode v40 2022.04, Ensembl 106). A background removal step was performed in all samples using CellBender^125^ with parameters --fpr 0.01 and --epochs 150. Full count matrices with background RNA removed were further analyzed using Seurat^126^. Cells with more than 10% of mitochondrial counts were eliminated and the number of genes detected was also used for filtering the data (see Extended Data Table 8 for specific parameters applied for each sample). Each object was normalized using Seurat’s LogNormalize method and transformed using ScaleData function removing the unwanted variation originated from mitochondrial contamination or the cell cycle stage. Upon examining Elbow plots, 20 Principal Components were considered significant for unsupervised clustering with the FindNeighbors and FindClusters Seurat functions. Cluster identities were assigned based on the genes specifically marking each cluster according to Seurat’s FindMarkers function and comparing them to lineage markers uncovered in human preimplantation datasets^32,45^. Seurat objects belonging to the nontarg-CARGO or the LTR5Hs-CARGO blastoids were merged and the LTR5Hs-CARGO object was downsampled to the same number of cells than the nontarg-CARGO object for comparison purposes. Multiple iterations of downsampling were performed with comparable results. We subset cells belonging to the epiblast or the neo-epiblast into a single group and performed differential gene expression analysis using DESeq2 on the sample-level aggregated counts (pseudobulk). Specifically, DESeq2 tested the effect of the repression of LTR5Hs elements (nontarg-CARGO v. LTR5Hs-CARGO) and using the PIP-seq replicate as a covariate. Only genes with FDR 5% and fold changes <-1 or >1 were considered statistically significant.

### Projection of PIP-seq transcriptomes into the human embryo reference datasets

To identify the human embryo counterparts of the transcriptomes of cells dissociated from nontarg-CARGO or LTR5Hs-CARGO blastoids, we projected such transcriptomes into a collection of human embryo reference datasets^29,32,45–48^. Counts from each gene in each cell (slot “counts” in Seurat) were extracted and uploaded to a human embryogenesis online prediction tool (https://petropoulos-lanner-labs.clintec.ki.se/)^49^. The identified annotations and UMAP values were used in our plots and conclusions.

### SMART-seq scRNA-seq data analysis including transposons

Raw data from Kagawa et al.^26^ was downloaded from the Gene Expression Omnibus database entry GSE177689. Smart-seq2 PCR adapters were trimmed using Skewer^118^ and the resulting reads were aligned using HISAT2^124^ with the parameters --dta --no-mixed --no-discordant -k 100, to allow enough multimappers for transposon analysis. The resulting bam files were processed with the scTE^127^ software for transposon family identification and quantification. The resulting matrix was subset to contain only cells of 96 h blastoids and it analyzed using Seurat, filtering cells with more than 25% of mitochondrial counts, less than 2000 or more than 16000 genes. Downstream unsupervised clustering was performed using 20 principal components. For comparing Kagawa et al. and this manuscript’s blastoids, we integrated a nontarg-CARGO scRNA-seq object with the Kagawa et al. data after removing the transposons using default Seurat data integration functions.

### Human-mouse and human-marmoset transcriptomes comparisons

LTR5Hs-regulated genes in the blastoids epiblast located within 250 kb of an LTR5Hs element were used as LTR5Hs target genes. To obtain genes expressed in the mouse epiblast we used a previously published table containing expression levels and orthology analysis of human, mouse and marmoset genes. Specifically, we used the epiblast data (early inner cell mass in the dataset)^61,128,129^. Only genes with an average expression of more than 2 FPKM were considered expressed. This cut-off was validated by visual inspection of scRNA-seq of mice Genes were assigned to evolutionary branches using data from the Gentree database^62^.

## Data availability

Datasets generated in this manuscript have been deposited in the Gene Expression Omnibus repository. CUT&RUN accession: GSE262191. Bulk RNA-seq accession: GSE296554. PIP-seq accession: GSE262329. ChIP-seq accession: GSE296555.

## Acknowledgements

We thank members of the Wysocka lab for their continued help and critical reading of the manuscript, especially to Saman Tabatabaee for technical help with recording the blastoids movie. We thank Dr. Fabian Suchy and Dr. Joydeep Bhadury for sharing technical expertise on human embryogenesis and Dr. Hideki Masaki for his help obtaining the hnPSCs. We also thank Dr. Conchi Estaras and Dr. Lina Colella for helpful discussions and Dr. Arttu Jolma and Dr. Tim Hughes for sharing unpublished data on ZNF729. The HERVKcon plasmid was a gift from Dr. Paul Bieniasz^40^, the pAAV-GFP plasmid a gift from Dr. John Gray (Addgene #32395), and the pDGM6 plasmid a gift from Dr. David Russell (Addgene #110660). R.F. was funded by an EMBO Long-term postdoctoral fellowship (ALT-1-2019) and a Cancer Research Institute-Bristol Myers Squibb fellowship. S.W. was supported by a Stanford Graduate Fellowship. J.W. was funded by HHMI, and a Lorry Lokey endowed professorship.

## Author contributions

R.F. and J.W conceptualized the study. R.F. designed, executed, and analyzed the experiments with assistance from S.W, O.C, and T.S. and supervision from J.W. R.F. and T.S performed statistical analyses. H.N. reviewed the study. The manuscript was written by R.F. and J.W. with contributions from other coauthors.

---

**Extended Data Figure 1.**
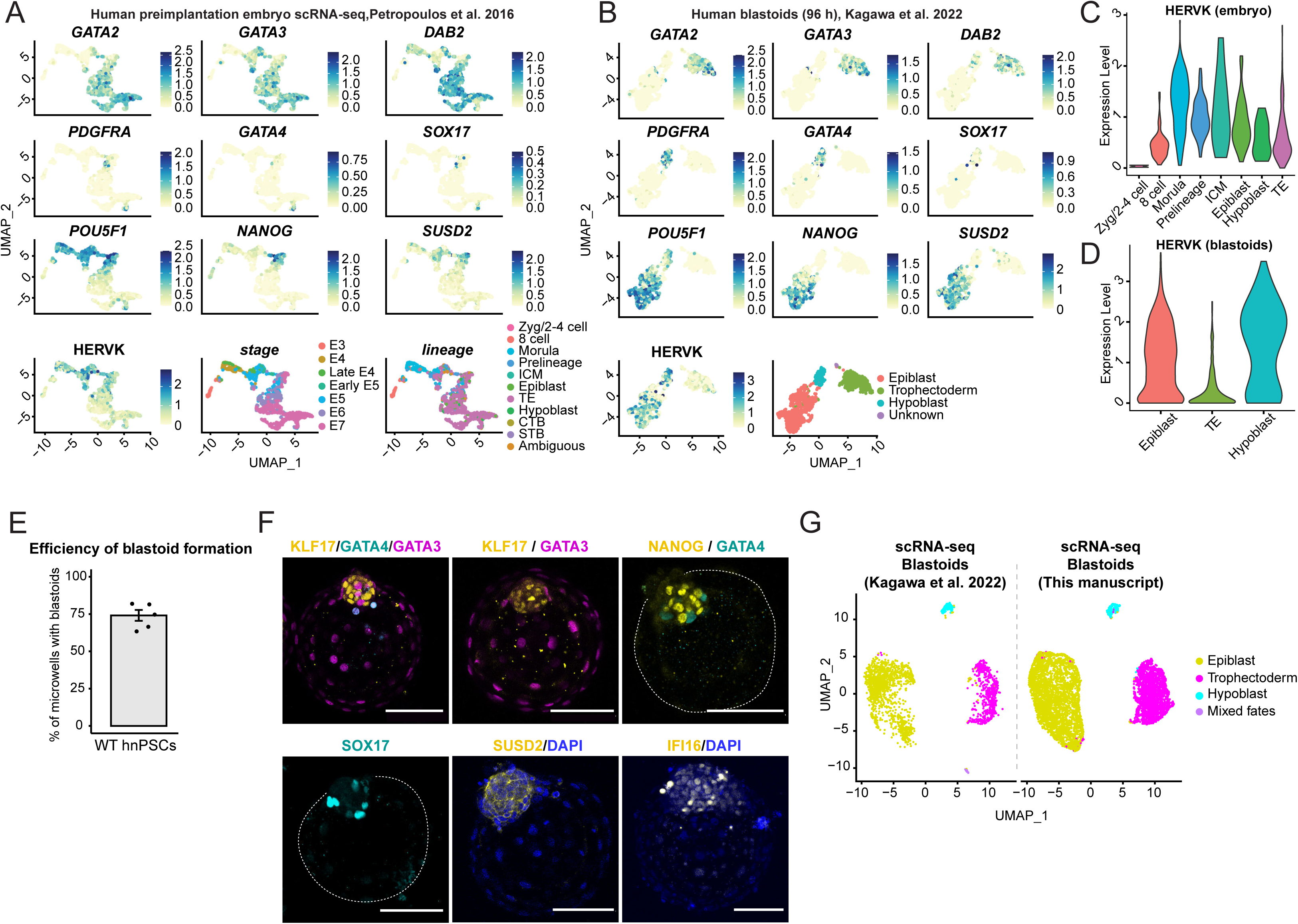
Benchmarking a human blastoid protocol to study HERVK in human preimplantation development. a. UMAP of expression of the indicated lineage markers and HERVK in human preimplantation single cells. Bottom right two panels display annotation of data using stage (the prefix E is equivalent to days) or lineages (data and annotations from 32,49). b. UMAP of expression of the indicated lineage markers and HERVK in single cells dissociated from blastoids at 96 h. Bottom right panels display annotation of data using lineages (data and annotations from26). c. and d. Violin plots of HERVK normalized expression levels at the different lineages present in human preimplantation embryos (c) or 96 h blastoids (d). e. Bar plot displays efficiency of blastoid formation with wild type hnPSCs, n=4 biologically independent replicates. f. Representative confocal images (n=3 or more) of immunostaining of blastoids derived from wild type hnPSCs with lineage-specific markers: KLF17, NANOG, SUSD2, IFI16 (yellow, epiblast), GATA4, SOX17 (cyan, hypoblast), and GATA3 (magenta, trophectoderm). White bar represents 100 um. g. UMAPs representing integrated single cell RNA-seq datasets from the previously reported human blastoid derivation protocol (Kagawa et al. 202226) and blastoids generated and benchmarked in this manuscript. Colors represent the blastoid lineages.

**Extended Data Figure 2.**
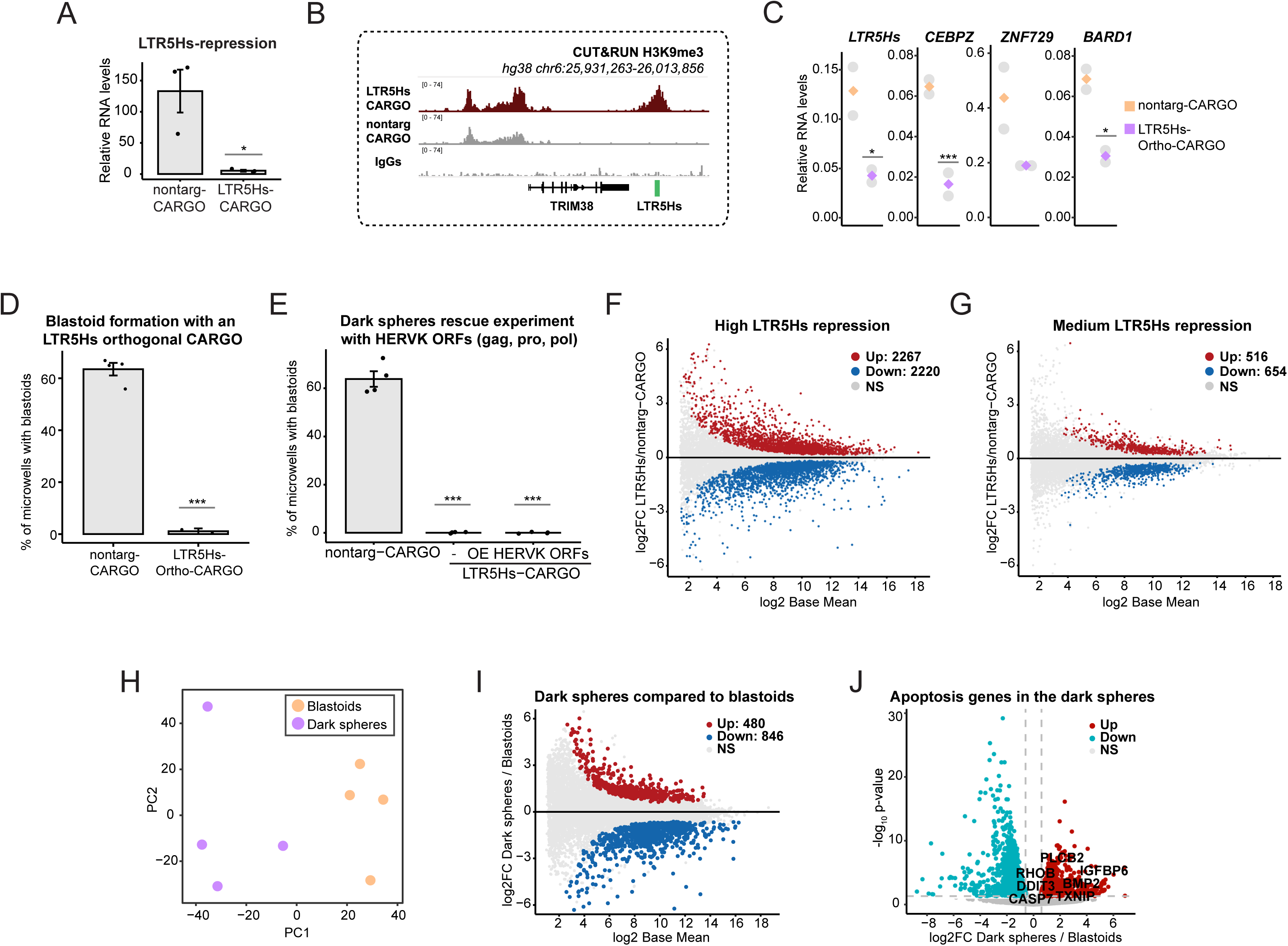
LTR5Hs-driven gene regulatory changes underlie blastoid formation potential. a. LTR5Hs-CARGO allows for efficient repression of LTR5Hs-originating transcripts in hnPSCs. RT-qPCR depicting LTR5Hs RNA levels measured with Taqman probes and normalized to the RPL13 gene in nontarg-CARGO and LTR5Hs-CARGO cells, n=3, unpaired two-tailed t-test, *<0.05. b. IGV genome browser capture of H3K9me3 CUT&RUN profiles of nontarg-, LTR5Hs-CARGO, and IgGs over the hg38 coordinates chr6:25931263-26013856. c. LTR5Hs-Ortho-CARGO allows for efficient repression of LTR5Hs-originating transcripts and LTR5Hs-regulated genes in hnPSCs. Plots depict RNA levels of indicated transcripts in nontarg-CARGO and LTR5Hs-Ortho-CARGO expressing hnPSCs, measured by conventional qPCR (n=2). Unpaired two-tailed test, *<0.05, ***<0.001. d. Bar plot representing blastoid formation efficiencies of nontarg-CARGO and LTR5Hs-Ortho-CARGO hnPSCs edited cell populations. KRAB-dCas9 is driven by an Ef1a promoter to ensure robust repression in individual cells (n=4 for nontarg-CARGO and n=2 for LTR5Hs-Ortho CARGO, two biological replicates). Unpaired two-tailed test, ***<0.001. e. Bar plots representing blastoid formation efficiencies of nontarg-CARGO hnPSCs clones (n=4), LTR5Hs-CARGO high repression clones (n=3) and LTR5Hs-CARGO high repression clones expressing a transgene encoding the HERVK proteins gag, pro, and pol (HERVKcon plasmid40 subcloned into piggyBac with the LTR substituted by an Ef1a promoter). f. MA plot representing bulk RNA-seq results of LTR5Hs-CARGO high repression clones (n=4 clones, in biological triplicates) compared to nontarg-CARGO clones (n=3). Y-axis shows gene expression fold changes in LTR5Hs-CARGO high repression vs nontarg-CARGO hnPSCs. FDR 5%. Numbers of significantly misregulated genes are indicated. g. MA plot representing bulk RNA-seq results of LTR5Hs-CARGO medium repression clones (n=5, in biological duplicates or triplicates) compared to nontarg-CARGO clones. Y-axis shows gene expression fold changes in LTR5Hs-CARGO medium repression vs nontarg-CARGO hnPSCs. FDR 5%. Numbers of significantly misregulated genes are indicated. h. Principal component analysis of the bulk RNA-seq transcriptomes obtained from blastoids and dark spheres obtained upon LTR5Hs-CARGO induction (n=4). i. MA plot representing bulk RNA-seq results of blastoids and dark spheres obtained upon LTR5Hs repression. Y-axis shows gene expression fold changes in dark spheres vs blastoids (n=4) FDR 5%. Numbers of significantly misregulated genes are indicated. j. Volcano plot of bulk RNA-seq of dark spheres compared to blastoids (n=4). Labeled dots represent statistically significant genes involved in apoptosis according to GSEA curated gene sets. FDR 5%.

**Extended Data Figure 3.**
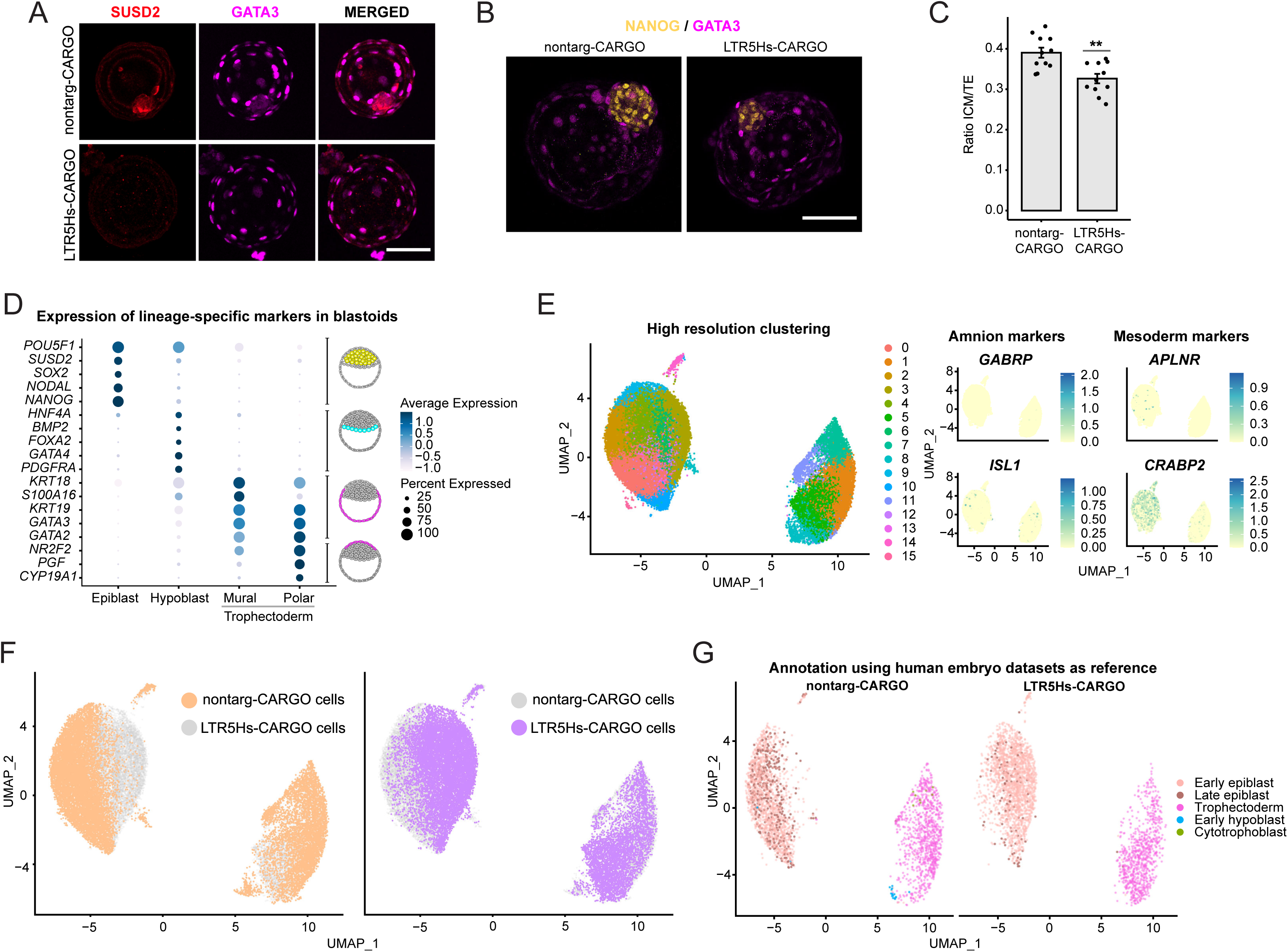
LTR5Hs-repression in human blastoids impairs lineage identity. a. Representative confocal images of a nontarg-CARGO and LTR5HS-CARGO blastoid immunostained with antibodies against the epiblast marker SUSD2 (red) and the trophectoderm marker GATA3 (magenta) n=3 blastoids from two independent biological replicates. White bar represents 100 um. b. Representative confocal images of a nontarg-CARGO and LTR5HS-CARGO blastoid immunostained with antibodies against the epiblast marker NANOG (yellow) and the trophectoderm marker GATA3 (magenta) n=3, white bar represents 100 um. c. Bar plots representing the ratio between the size of the inner cell mass (ICM) and the size of the trophectoderm cavity (TE) in nontarg-CARGO and LTR5Hs-CARGO blastoids. Each dot represents measures from tens of blastoids in each independent biological replicate (n=11, 7 biological replicates but 4 of them with technical replicates). Unpaired t-test, **<0.01. d. Dot plot showing the expression of genes indicated on the y-axis in cells assigned to the clusters indicated on the x-axis. Cartoons on the right highlight the location of cells expressing the subset of genes in the human blastocysts/blastoids. e. High resolution clustering (resolution = 1) of PIP-seq results (left) and UMAP of the expression of amnion markers and mesoderm markers (right) in those clusters. f. UMAP of transcriptomes from single cells dissociated from nontarg-CARGO blastoids (left, orange) and LTR5Hs-CARGO blastoids (right, purple) coloring uniquely cells from each origin for better visualization of the UMAP depicted in Figure 2E. g. UMAP representing the nontarg-CARGO and LTR5Hs-CARGO transcriptomes colored by the assigned human embryo transcriptomic counterpart using the reference datasets^29,32,45–48^ using a computational tool published in^49^.

**Extended Data Figure 4.**
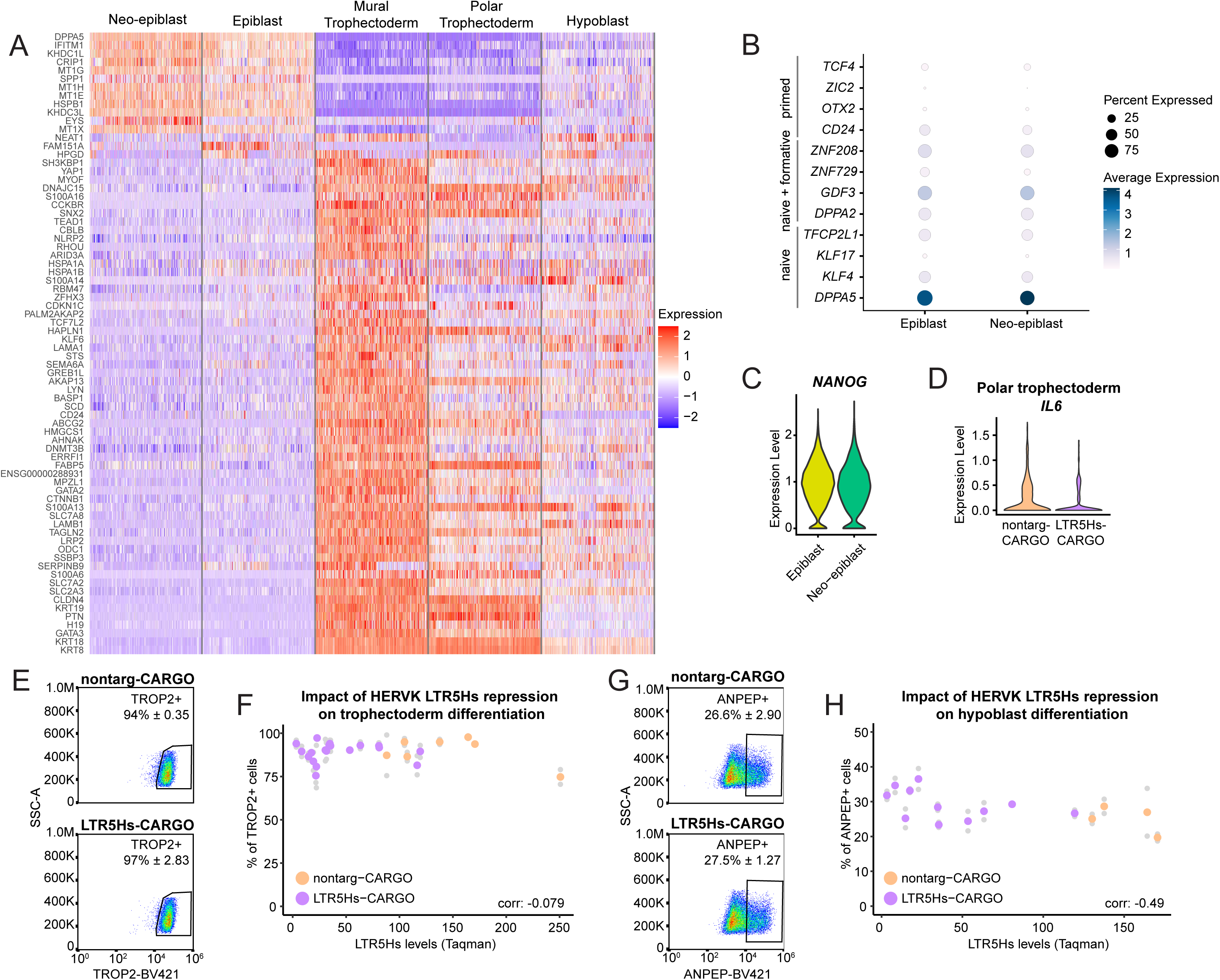
Lineage identity features of LTR5Hs-repressed blastoids. a. Heatmap representing the expression of genes identified as markers of the clusters indicated on top in the PIP-seq data. Color scale represents z-score. b. Dot plot showing the expression of naïve, formative and primed pluripotency genes indicated on the y-axis in cells assigned to the epiblast or neo-epiblast clusters indicated on the x-axis. c. Violin plot of NANOG normalized expression levels in the epiblast and neo-epiblast clusters. d. Violin plot of IL6 normalized expression levels in nontarg-CARGO and LTR5Hs-CARGO blastoids. e. Flow cytometry density plots representing the analysis of TROP2+ cells compared to size scatter in 2D cultures of nontarg-CARGO (7 clonal cell lines in biological duplicates) and LTR5Hs-CARGO hnPSCs (15 clonal cell lines in biological duplicates) upon 3 days of differentiation using the trophectoderm protocol described in50. f. Representation of the quantitative results of the flow cytometry experiments described in e. Correlation represents Pearson’s. f. Flow cytometry density plots representing the analysis of ANPEP+ cells compared to size scatter in 2D cultures of nontarg-CARGO (4 clonal cell lines in biological duplicates) and LTR5Hs-CARGO hnPSCs (11 clonal cell lines in biological duplicates) upon 3 days of differentiation using the hypoblast protocol described in51. h. Representation of the quantitative results of the flow cytometry experiments described in g. Correlation represents Pearson’s.

**Extended Data Figure 5.**
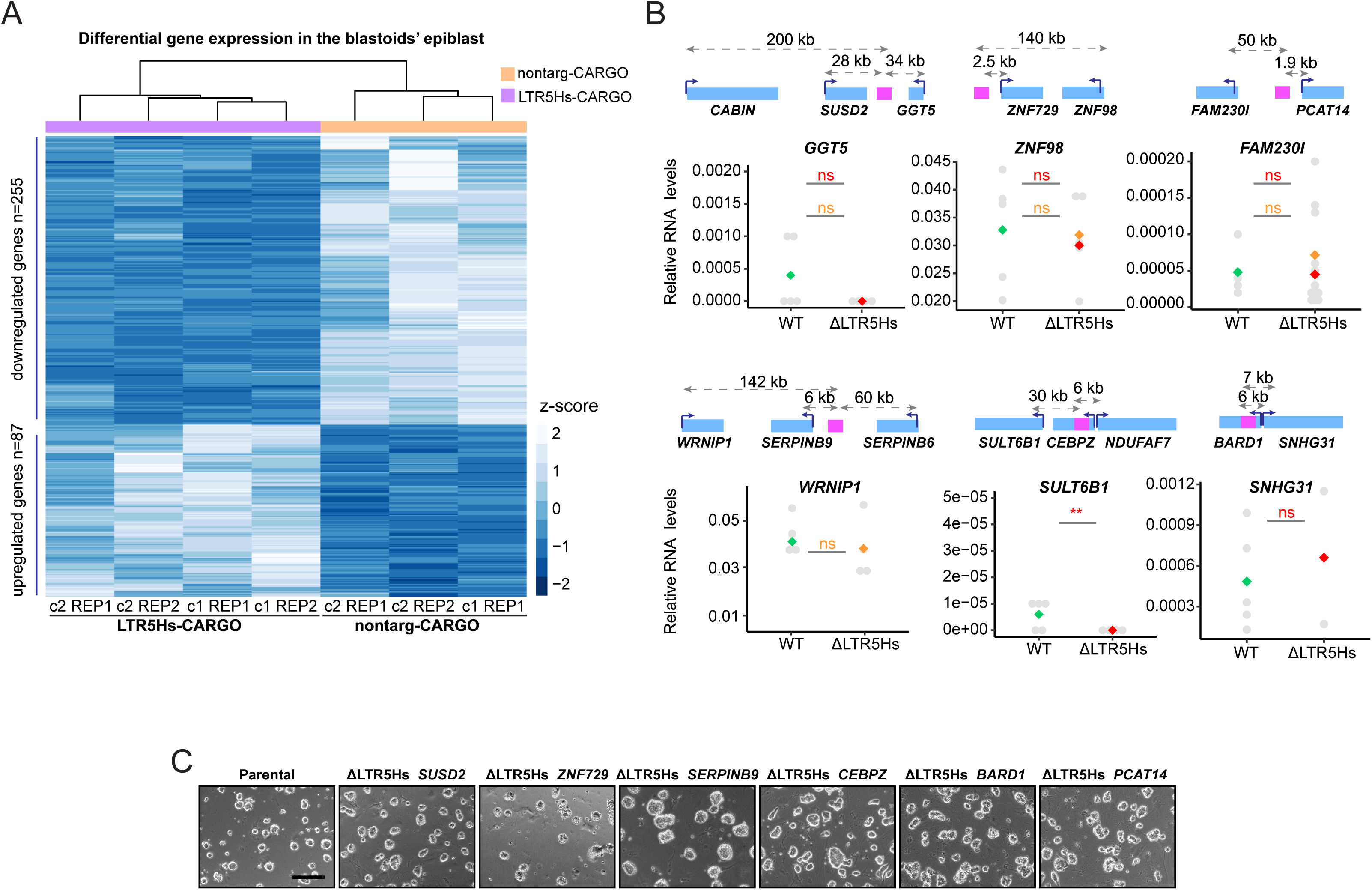
LTR5Hs regulates gene expression in the epiblast of blastoids. a. Heatmap and hierarchical clustering representing differentially expressed genes in the blastoids generated from nontarg-CARGO or LTR5Hs-CARGO hnPSCs cell lines (two clonal cell lines from each condition, n=2 PIP-seq biological replicates per clone, except for nontarg-CARGO clone 1, n=1). Color scale represents z-score. b. RT-qPCR results of genes unaffected by LTR5Hs deletions at gene loci shown in Figure 3E. Plots display the expression of the indicated genes in wild type or ΔLTR5Hs hnPSCs. RNA values are normalized relative to RPL13A. Above each plot, schematic depiction of the locus is shown. Blue rectangles indicate genes, pink indicates the closest LTR5Hs element, the number on top of the dashed arrows displays distance from the promoter to the LTR5Hs. Grey dots represent expression values obtained in each clone, green, orange, and red dots represent median values. Unpaired two-tailed test, ***<0.001, **<0.01, *<0.05, asterisk color indicates p-value calculated for the homozygous (red) or heterozygous (yellow) clones. ns: not significant. c. Representative bright field images of Δ LTR5Hs hnPSCs with the genes affected by each deletion indicated above each image. Black bar represents 200 um.

**Extended Data Figure 6.**
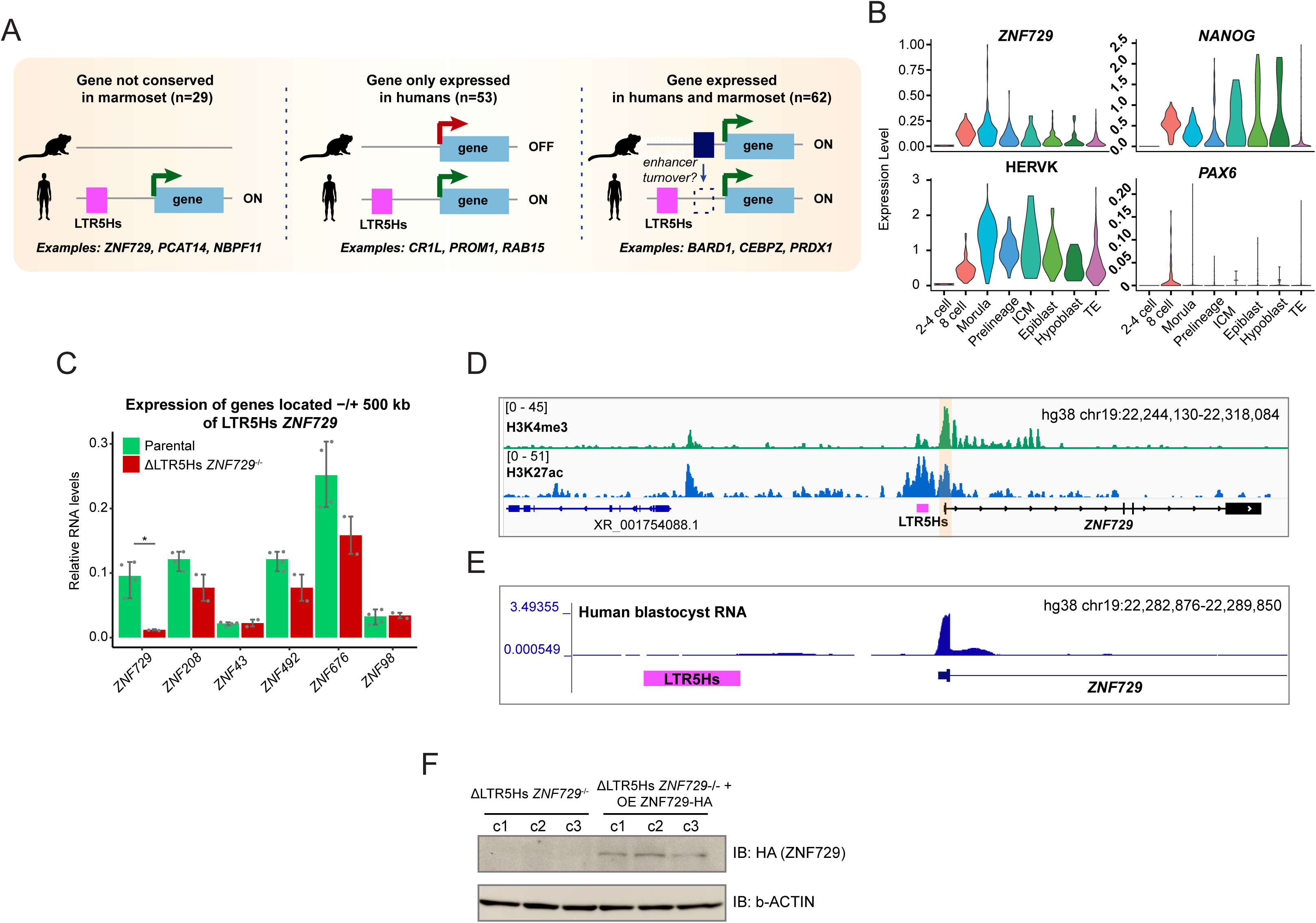
Expanded characterization of the LTR5Hs diversification of the transcriptome and ΔLTR5Hs ZNF729 hnPSCs. a. Expression and conservation of the LTR5Hs-regulated genes in marmosets. Human epiblast genes regulated by LTR5Hs in cis in blastoids were divided into three groups, based on the status of expression in the marmoset’s epiblast and conservation of the gene itself using data published in 61. The number of genes in each of the three groups is indicated on top, examples of genes within each group are shown at the bottom. b. Violin plots of ZNF729 and HERVK normalized expression levels in annotated lineages of human preimplantation scRNA-seq data32. NANOG and PAX6 are depicted as a reference for a highly or lowly expressed transcription factor, respectively. c. RT-qPCR expression analysis of genes locates within ∼1Mb of the LTR5Hs ZNF729 locus. Plot displays the expression of the indicated genes in parental (green, n=4) or ΔLTR5Hs ZNF729-/-(red, n=2) hnPSCs. RNA values are normalized relative to RPL13A. Unpaired two-tailed test, *<0.05. d. IGV browser capture showing ChIP-seq signal of H3K4me3 (top, green) and H3K27ac (bottom, blue) at the LTR5Hs ZNF729 locus, hg38 chr19:22,244,130-22,318,084. e. UCSC browser capture of isoform resolved transcriptomic data in human blastocysts70 at the LTR5Hs ZNF729 locus (hg38 chr19: 22282876-22289850). f. Western blot validating the expression of the ZNF729 protein from the integrated ZNF729-HA cDNA transgene in three clonal cell lines of ΔLTR5Hs ZNF729 hnPSCs. Membrane was blotted with antibodies against the HA tag and β-ACTIN as loading control blotted on the same membrane. For gel source data, see Extended Data Figure 9.

**Extended Data Figure 7.**
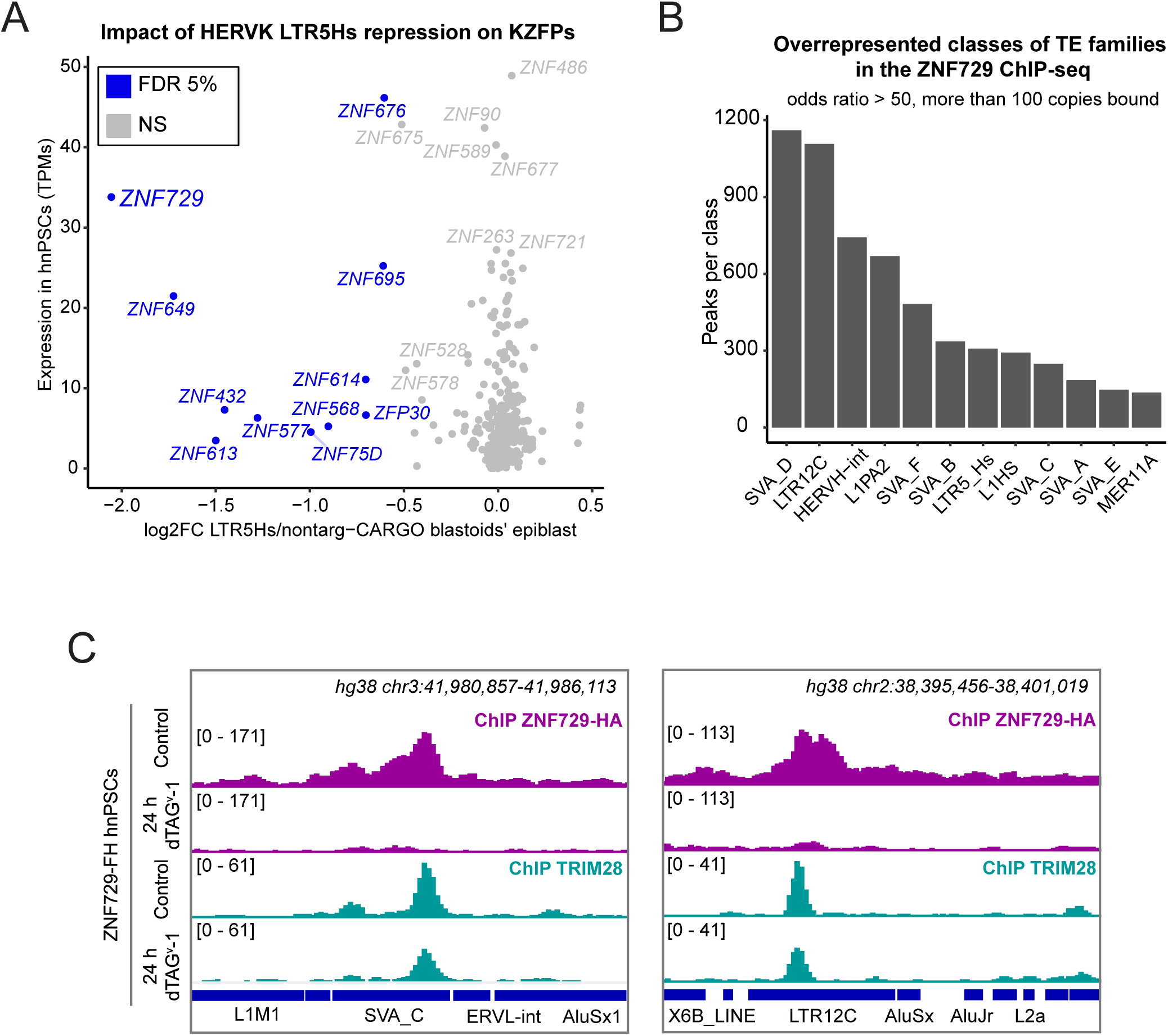
ZNF729 binds transposable elements in hnPSCs. a. ZNF729 is the most dysregulated KZFP in the blastoids’ epiblast upon LTR5Hs-repression. Scatter plot depicting expression changes in KZFPs (curated in Krabopedia132) in the blastoids’ epiblast upon LTR5Hs repression (X axis) vs their expression in hnPSCs (Y axis). FDR 5%. b. Histogram representing overrepresented classes of TEs in the ZNF729-FH bound DNA regions. Only the top 12 families are represented. Y-axis displays the number of bound TEs from each family. c. IGV genome browser capture of ZNF729-FH ChIP-seq signal (top two tracks, purple) or TRIM28/KAP1 ChIP-seq signal (bottom two tracks, turquoise) in DMSO treated (control) and 24 h dTAGv-1 treated ZNF729-FH at loci containing TEs bound by ZNF729 (SVA_C, left panel, hg38 chr3:41980857-41986113; LTR12C, right panel hg38 chr2:38395456-38401019).

**Extended Data Figure 8.**
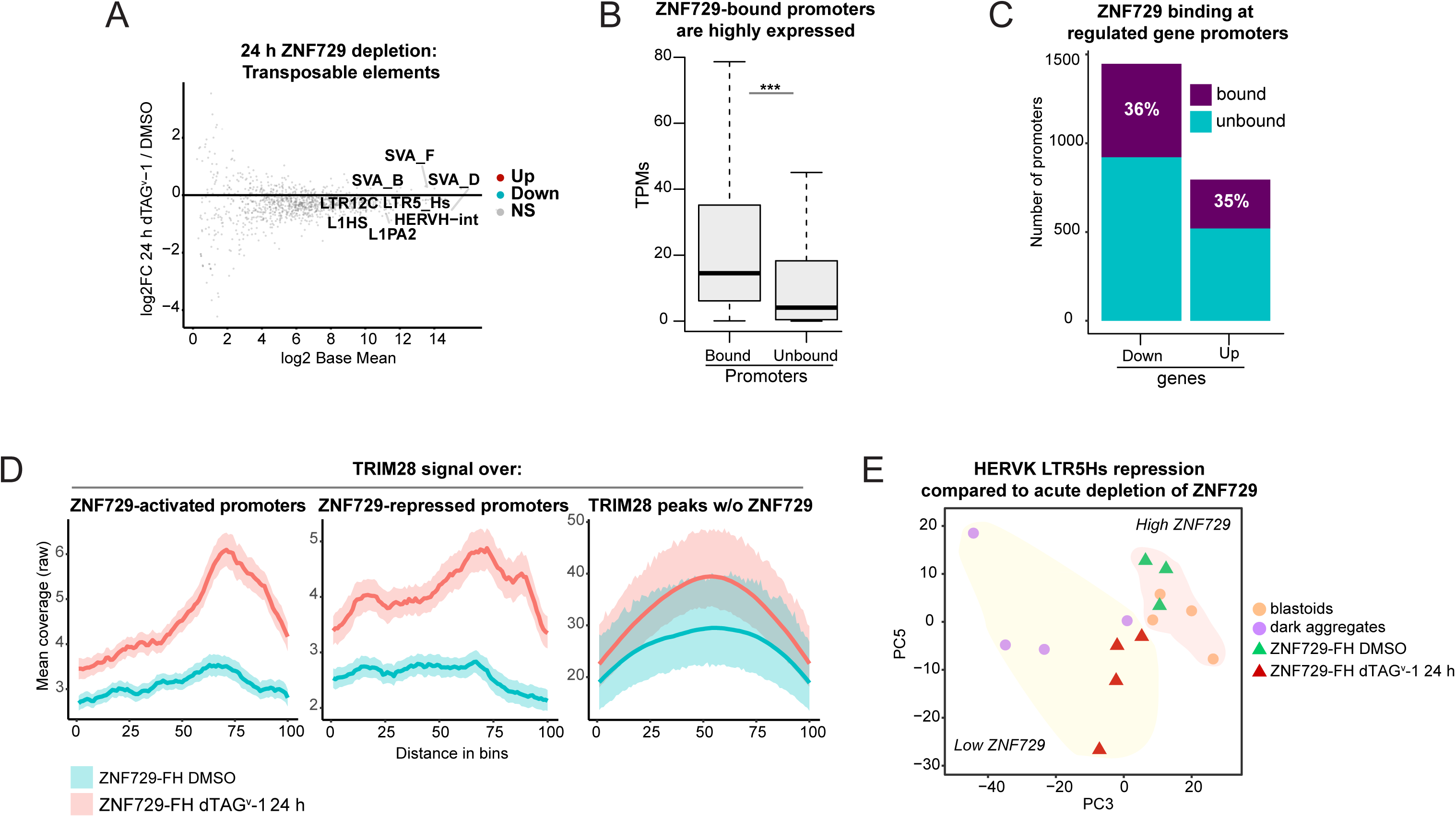
ZNF729 directly exerts activator and repressor function. a. Low impact of ZNF729-FH depletion on the expression of TEs. MA plot represents TEtranscript81-obtained data quantified with DESeq252. FDR 5%. Top ZNF729-bound TE families are labeled. b. Expression from gene promoters bound or unbound by ZNF729. Box plots show basal hnPSC expression (in TPMs) of genes bound or unbound by ZNF729, Wilcoxon test, *<0.05. c. Stacked bar plot represents number of down- or upregulated genes upon ZNF729-FH depletion for 24 h with dTAGv-1 and the percentage of them directly bound by ZNF729 at their promoters (−1kb/200bp around the TSS). d. Metagene plots showing TRIM28 ChIP-seq signal over promoters activated or repressed by ZNF729 and over a negative control (TRIM28 bound regions not overlapping with ZNF729). Green represents ZNF729FH DMSO (control) and red dTAGv-1 treated. e. PCA analysis of bulk transcrip-tomes form blastoids, dark spheres formed upon high LTR5Hs repression, and DMSO-treated or dTAGv-1 treated ZNF729-FH hnPSCs. Red shade represents “normal / high” levels of ZNF729 (blastoids, and DMSO treated ZNF729-FH cells). Yellow shade represents “low” ZNF729 levels (dark spheres with high LTR5Hs repression, and dTAGv-1 treated ZNF729-FH hnPSCs). Note that high ZNF729 and low ZNF729 samples separate along PC3.

**Extended Data Figure 9.**
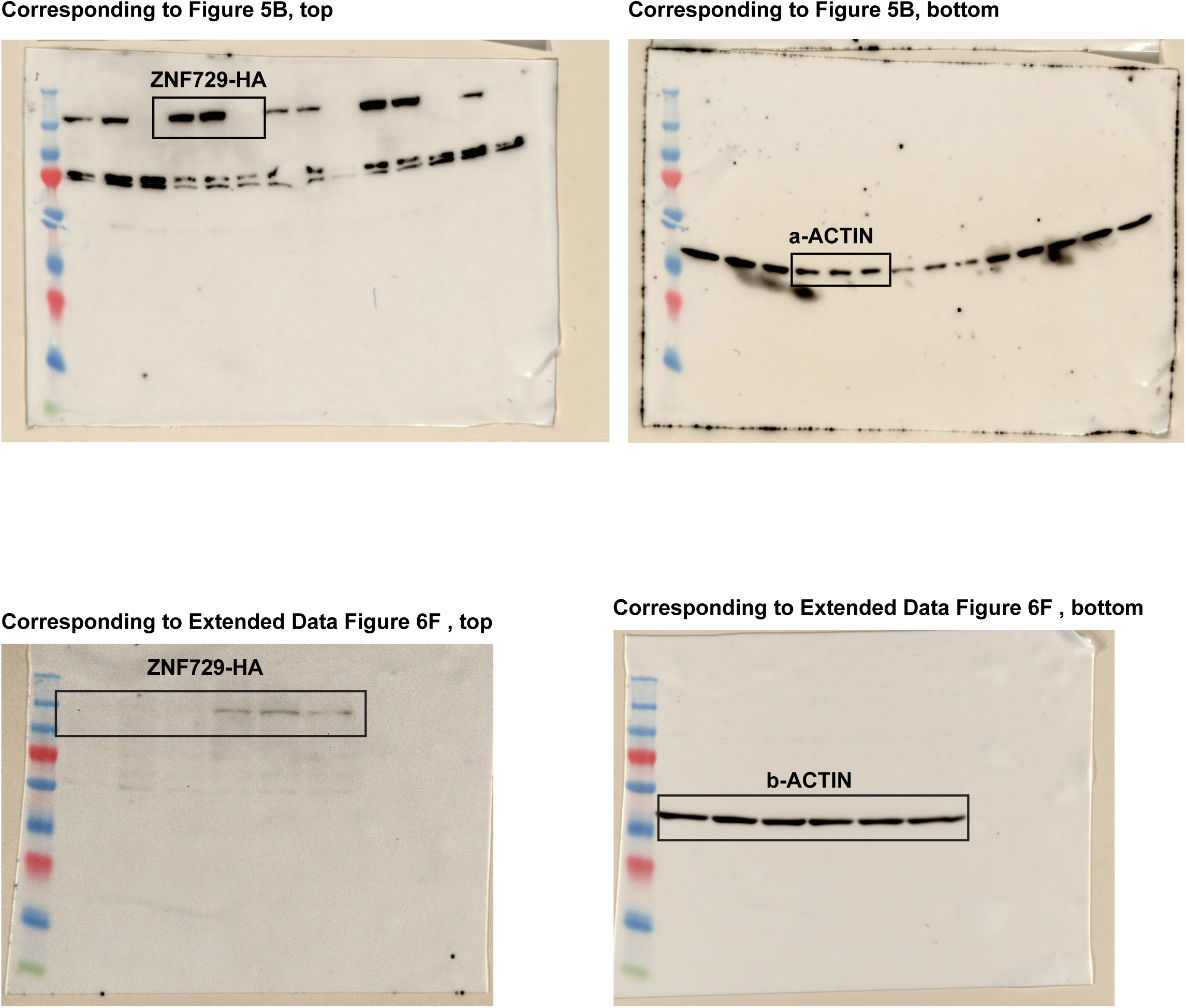
Uncropped western blot images.

